# Distinct allosteric remodeling of HIV-1 Env dynamics on virions by gp41-directed antibodies reveals two modes of neutralization

**DOI:** 10.64898/2026.01.27.702099

**Authors:** Wang Xu, Narendra Kumar Gonepudi, Junyu Liu, Yufan He, Revansiddha Katte, Ran Wang, Harry Baffour Awuah, Yang Han, Baoshan Zhang, Jian Yu, Bo Hu, David D. Ho, Priyamvada Acharya, Peter D. Kwong, Maolin Lu

## Abstract

HIV-1 envelope glycoprotein (Env), a gp120–gp41 trimer, undergoes coordinated conformational changes that drive membrane fusion and allow immune evasion by transiently concealing neutralization-sensitive epitopes. Most broadly neutralizing antibodies (bNAbs) target gp120, whereas a distinct subset recognizes conserved gp41 regions, such as the fusion peptide and the membrane-proximal external region; however, their impact on Env dynamics and associated neutralization mechanisms remains unclear. Using bioorthogonal tagging for single-molecule FRET, we monitored real-time bNAb-induced conformational sampling of Env on intact virions. Most gp41-directed bNAbs allosterically stabilized the prefusion-closed (PC) state, whereas the bivalent 10E8.4/iMab favored both PC and CD4-bound open (predominant) states. Antibodies redistributed the conformational populations of Env with modest kinetic effects, preserving the sequential transition pathway. These findings reveal two modes of neutralization for gp41-directed antibodies, fixing the prefusion-closed conformation and opening it up – in both cases, with neutralization occurring via long-range allosteric control of Env dynamics.

## Introduction

The HIV-1 envelope glycoprotein (Env) mediates the fusion of the viral and host cell membranes, enabling viral entry.^1–3^ As the sole viral protein coated on the virion surface, Env is a central focus for vaccine design and the development of therapeutic antibodies, small-molecule inhibitors, and fusion blockers.^3,4^ Env assembles as a trimer of noncovalently associated gp120–gp41 heterodimers, with gp120 engaging the receptors CD4 and coreceptor (CCR5 or CXCR4) and gp41 driving membrane fusion.^2,3,5–8^ These Env trimers are present at low copy number (roughly 5–15 per virion), and although their exact stoichiometry remains unresolved, productive fusion may require the coordinated action of multiple trimers.^9–16^ Env is highly dynamic, undergoing large conformational rearrangements.^17–23^ A key early step is the sequential opening of the three gp120 protomers, which swing outward to form a fully open, CD4-bound trimer that exposes the CD4- and coreceptor-binding sites and primes gp41 for fusion activation.^24–26^ This is followed by transient intermediate states and the refolding of gp41 into a post-fusion conformation that leads to membrane fusion.^2,3,27–31^ This conformational plasticity, combined with a dense glycan shield and rapid sequence diversification, enables Env to evade immune pressure by transiently concealing or exposing neutralization-sensitive epitopes, known as “conformational masking.”^3,32,33^ Despite these defenses, a distinct subset of antibodies, known as broadly neutralizing antibodies (bNAbs), can neutralize diverse HIV-1 isolates by targeting conserved sites of vulnerability across Env.^4,33,34^ These epitopes span multiple structural regions, including the V1V2 apex, V3 glycan supersite, CD4-binding site, gp120–gp41 interface, fusion peptide (FP), and the membrane-proximal external region (MPER).^4,33,34^ Eliciting potent Env-directed bNAbs remains a central goal of HIV-1 vaccine development.^4,33–36^ Understanding how Env dynamics, intrinsically linked to its fusogenic function, shapes bNAb-mediated neutralization is important for guiding antibody-based therapies and designing next-generation immunogens.

HIV-1 neutralization by bNAbs is multifaceted and likely involves overlapping mechanisms.^4,33,37^ By exploiting structural and functional constraints on Env, bNAbs can neutralize Env by competing with receptor engagement, disrupting the conformational changes required for membrane fusion, penetrating into Env glycan holes, and/or trapping the trimer into specific states. Most current insights derive from static cryo-EM and crystal structures of soluble Env constructs,^4,35,36^ which capture discrete conformations with atomic details of molecular interactions but do not reveal the sequence, timing, or rates of transitions between states. For gp41 epitopes, specifically FP and MPER, which are recognized by some of the most potent bNAbs, it is unclear whether antibody binding exerts long-range allosteric effects on native Env dynamics, and the impact on the distribution and kinetics of conformational sampling along the gp120–gp41 axis remains poorly defined. Among those, the highly potent 10E8.4/iMab,^38,39^ which combines an MPER-directed arm with the CD4-targeting antibody Ibalizumab (iMab) and is currently in clinical trials, represents a particularly promising therapeutic strategy. However, neither structural nor dynamic analyses of its complex with Env have yet been performed, leaving unresolved both the Env conformations targeted by 10E8.4/iMab and its effects on Env dynamics.

Here, we address these knowledge gaps by combining site-specific bioorthogonal tagging^40–42^ with single-molecule Förster Resonance Energy Transfer (smFRET)^43–46^ to visualize the real-time conformational dynamics of full-length Env on intact virions. While prior work has analyzed gp120-directed antibodies using smFRET with fluorophores labeled on gp120,^17,27^ gp41-directed antibodies have never been systematically investigated by smFRET, in part due to the lack of a suitable FRET-pair labeling system. By using dual click-labeling at gp120–gp41 positions, here we characterize the conformational ensemble of native Env and define how bNAbs targeting the gp120–gp41 interface, FP, and MPER allosterically remodel trimer dynamics. Our results reveal previously unobserved shifts in Env conformational sampling and two dynamic modes of neutralization, accompanied by modest changes in transition kinetics that preserve the intrinsic opening pathway. These findings provide a unique perspective on how gp41-directed bNAbs alter Env conformational dynamics through long-range allosteric effects, offering mechanistic insight into HIV-1 neutralization and informing strategies to exploit these vulnerabilities for vaccine and therapeutics development.

## Results

### Dual amber-click labeling and functional validation of Env trimers for smFRET

To investigate the conformational dynamics of full-length Env on intact HIV-1 virions, we employed a minimally invasive amber-click labeling strategy that combines amber suppression with click chemistry for smFRET studies (**Figs. S1–S2**). Our previously developed amber-free HIV-1 system eliminates background amber (TAG) stop codons, allowing efficient site-specific incorporation of unnatural amino acids (ncAAs) into the target HIV-1 protein of interest.^42,47^ By using this platform, we introduced TAG codons at S401^TAG^ in gp120 and R542^TAG^ in gp41 of Env_BG505_ (a transmitted/founder primary isolate, **Fig. 1A**) on the HIV-1_Q23_ viral backbone (HIV-1_Q23_ Env_BG505_). These tagging sites (**Fig. S1**) were strategically selected along the gp120–gp41 axis based on available structural details,^6,21,48,49^ allowing them to report on inter-subunit movements during conformational transitions. Through genetic code expansion with an orthogonal tRNA/tRNA synthetase pair (tRNA^Pyl^/NESPylRS^AF^),^41,50^ we then incorporated the ncAA trans-cyclooct-2-en-L-lysine (TCO*A) at these amber-tagged sites for fluorophore attachment (**Fig. S2**).^47,51,52^ Site-specific conjugation of LD555-TTZ (donor, LD555: Cy3 derivative; TTZ: tetrazine) and LD655-TTZ (acceptor, LD655: Cy5 derivative) fluorophores was further achieved by click chemistry via strain-promoted inverse electron-demand Diels–Alder cycloaddition^53,54^ (SPIEDAC, **Fig. S2**).

**Figure 1.**
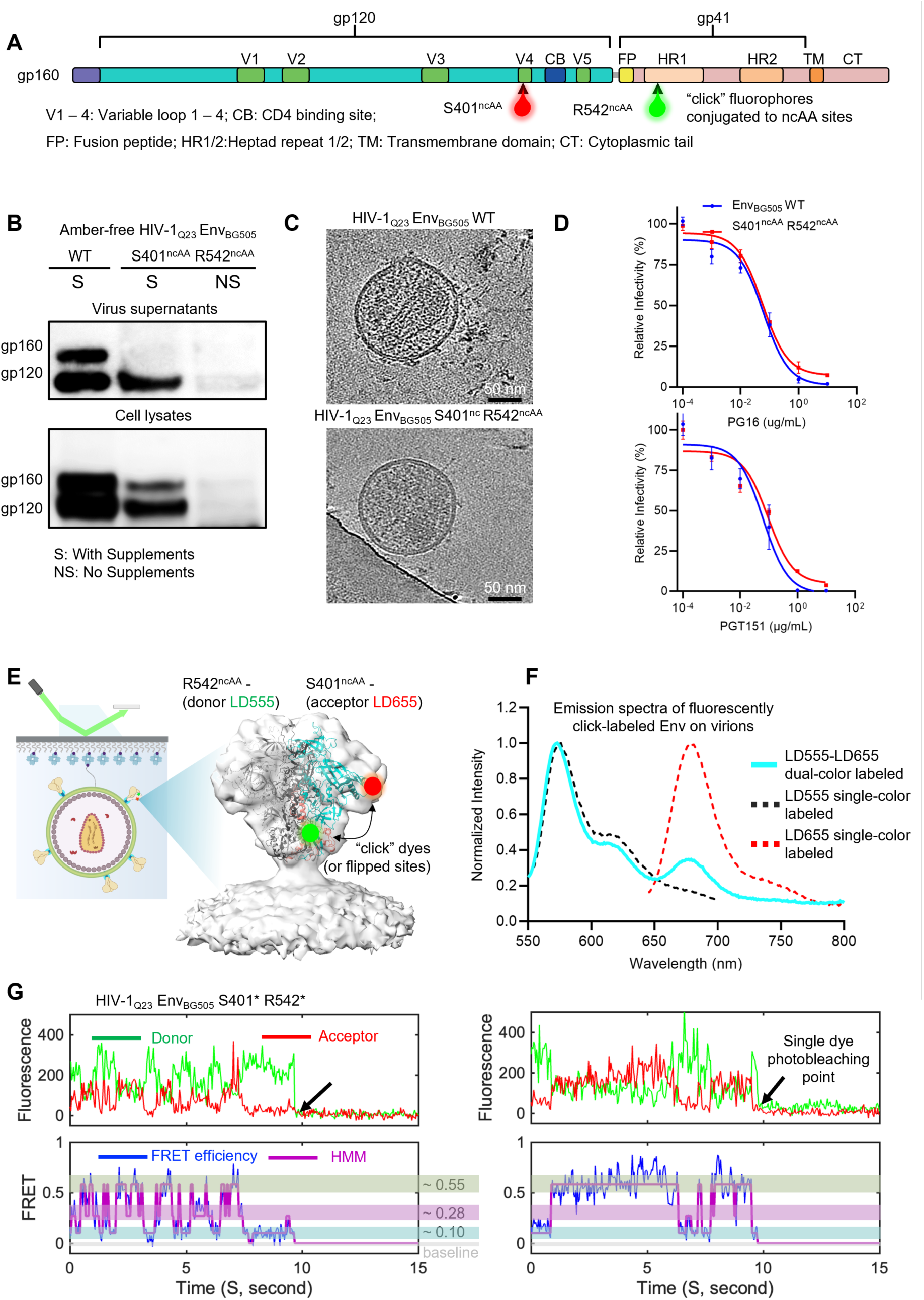
smFRET imaging of full-length Env on HIV-1 virions from the gp120–gp41 structural perspective. (**A**) Domain organization of BG505 Env with click fluorophores conjugated to ncAA sites at S401 on gp120 and R542 on gp41. “ncAA” refers to unnatural amino acids incorporated at amber (TAG) sites via amber suppression, and “click” dyes are site-specifically conjugated through click chemistry (see Fig. S2 for details). **(B – D)** Functional validation of HIV-1_Q23_ Env_BG505_ S401^ncAA^ R542^ncAA^ virions for smFRET studies. Immunoblotting (**B**), tomographic slices (**C**), and neutralization curves (**D**) confirm that the dual-ncAA incorporation does not compromise Env trimer functionality on virions used for smFRET. Viruses were generated by transfection with 100% of the indicated HIV-1 constructs, and the data shown represent the resulting particles. (**E**) HIV-1_Q23_ virions carrying fluorescently labeled full-length Env_BG505_, imaged by prism-based TIRF microscopy. The Env trimer (PDB 4ZMJ) was fitted into the electron density map of the membrane-bound Env trimer (EMD-21412). Two click fluorophores (donor LD555-TTZ and acceptor LD655-TTZ) were site-specifically introduced at S401^ncAA^ and R542^ncAA^ of a single protomer (gp120 in cyan, gp41 in pink; wild-type protomers in grey). Labeling sites are shown as green and red spheres, respectively. (**F**) Fluorescence emission spectra of HIV-1_Q23_ virions carrying click-labeled Env_BG505_ S401^ncAA^ R542^ncAA^, showing LD555 and LD655 signals from dual-color virions (solid cyan) compared with spectra from single-color–labeled virions (dashed lines). (**G**) Representative donor (green) and acceptor (red) fluorescence traces and corresponding FRET efficiency trajectories (blue) with hidden Markov modeling (HMM) idealization (magenta), from two individual ligand-free HIV-1_Q23_ virions carrying Env_BG505_ S401* R542* (left and right panels). The black arrow marks single-step photobleaching. Three FRET-populated states are indicated by color-coded bands.

Throughout this study, S401^TAG^ R542^TAG^ denotes the construct, S401^ncAA^ R542^ncAA^ denotes ncAA-incorporated but unlabeled virions, S401* R542* denotes their fluorescently labeled counterparts, and we refer to these sites as the gp120–gp41 (S401– R542) probe pair; the gp120 V1V4 (N136–S401) probe pair is used as a reference perspective.

To ensure that the tagged Env retained its functionality and was suitable for smFRET studies, we first validated dual-ncAA-incorporated Env (S401^ncAA^ R542^ncAA^) on intact virions as well as fluorescently labeled forms (S401* R542*). We obtained approximately 20% dual-amber suppression efficiency for HIV-1_Q23_ Env_BG505_ S401^TAG^ R542^TAG^, as determined by the relative infectivity of produced virions compared to wild-type (WT) Env (**Fig. S3A**). Importantly, click-chemistry labeling did not further reduce infectivity, with labeled and unlabeled virions exhibiting comparable infectivity (**Fig. S3B**), indicating that fluorescent labeling did not appreciably impair Env function.

The reduced infectivity relative to WT primarily reflects the inherent limitations of amber suppression in mammalian cells rather than loss of Env function, as discussed previously.^42,47,55,56^ Amber suppression must compete with endogenous translation termination, and unintended readthrough of endogenous or viral amber codons can interfere with protein expression and virus production. We minimized the latter by using an amber-free provirus.^47^ Dual-ncAA incorporation requires two successful suppression events within the same Env molecule, further reducing overall efficiency. Given these constraints, the ~20% dual-amber suppression efficiency achieved here is within the expected range and comparable to previous studies.^41,47^

Immunoblotting of prepared virions that were harvested from transfected cell supernatants confirmed that both TAG sites were successfully suppressed, proteolytically processed, and incorporated into amber-free HIV-1_Q23_ virions, with no detectable Env in non-suppression controls (**Fig. 1B**). Viral particle shape and size distributions were similar to wildtype, as revealed by cryo-ET, negative staining, and nanoparticle tracking analysis (**Fig. 1C**, **Fig. S3C–D**). A quantitative analysis of capsid morphology, Env spike incorporation, and the proportion of immature particles would be more informative. However, we reason that such analyses would require a substantially larger cryoET dataset, which is beyond the scope of the present smFRET-focused study of Env conformational dynamics.

The ncAA-incorporated virions also exhibited neutralization sensitivities comparable to WT when tested against trimer-specific bNAbs, including PG16 (apex-directed)^57^ and PGT151 (gp120–gp41 interface-directed)^58^ (**Fig. 1D**). Of note, these validations were performed using virions generated entirely from dual-amber–tagged Env constructs. Together, these results (**Fig. 1B–D**, **Fig. S3**) confirm proper Env expression, particle size, and preserved antigenicity, demonstrating that ncAA-incorporated Env is properly presented on intact virions and retains functional competence, and that click labeling does not affect virion infectivity. After functional validations and prior to single-molecule studies (**Fig. 1E**), we performed ensemble-level spectral characterization. Dually labeled virions with donor and acceptor fluorophores exhibited well-separated excitation and emission profiles, and single-color–labeled control virions displayed the expected single-emission signatures consistent with those of free dyes, confirming specific fluorophore conjugation (**Fig. 1F**, **Fig. S4**).

### Real-time visualization of global CD4-associated opening of native virus Env from two structural perspectives

We next performed smFRET imaging of fluorescently labeled Env trimers on intact virions using a lab-customized prism-based TIRF microscope (**Fig. 1E**). To optimize conditions for monitoring single FRET-labeled protomers, virions were generated by co-transfecting amber-free wild-type and S401^TAG^ R542^TAG^ Env constructs at a ratio based on the suppression efficiency. Under these conditions, most virions remained wild type and thus unlabeled following fluorophore conjugation. Post-labeled virions were immobilized on streptavidin-coated quartz slides via biotin–streptavidin interactions in a quartz–coverslip flow cell. Among labeled particles, those carrying a single gp120 bearing LD555/LD655 pairs within an otherwise wild-type Env background could be identified and analyzed based on their FRET signals. Thus, smFRET imaging selectively detected virions with a single dual-labeled protomer among mostly wild-type Env trimers, of which similar experimental designs (see **Methods** for details) have been successfully applied in prior smFRET studies of Env and other viral spike proteins.^17,27,59–61^

We analyzed hundreds of single-molecule trajectories that displayed anti-correlated donor and acceptor fluorescence intensity fluctuations, along with discrete single-step photobleaching events, as exemplified in **Fig. 1G** (fluorescence, top panels) for ligand-free S401* R542*. These features confirmed that individual traces correspond to single Env trimers in motion. The resulting FRET efficiency trajectories (**Fig. 1G**, bottom panels; FRET = FRET efficiency) report real-time conformational changes of individual Env trimers over their observable time period, until photobleaching occurred. Hidden Markov modeling (HMM) analysis^62,63^ of these trajectories, which deconvoluted each into the most probable sequence of discrete FRET states, consistently resolved three well-separated FRET populations (**Fig. 1G**): a low-FRET state with a mean value of ~0.1, an intermediate-FRET state with a mean of ~0.3, and a high-FRET state with a mean of ~0.55. This three-state model aligns well with our previous work and that of others.^17,19,24,47,64,65^

We assigned the newly observed FRET populations from the gp120–gp41 (S401–R542) axis, referencing the well-characterized states from the gp120 V1–V4 (N136–S401) axis (**Fig. 2**, **Figs. S5–6**, **Table S1**). Previous smFRET studies^17,19,24,47,64,65^ using the referenced axis have shown that native Env on virions predominantly resides in the pre-triggered (PT) state, transitions through the prefusion closed (PC) state, and fully opens into the CD4-bound open (CO) state (simplified in **Fig. 2A–B**, see **Fig. S5** for details). Because FRET values depend on donor–acceptor distance and probe geometry, we mapped the probe sites onto closed and open Env trimers (**Fig. 2C–D**) and next compared gp120–gp41 and gp120 V1–V4 profiles under matched conditions. This enabled confident assignment of FRET populations to PT, PC, and CO.

**Figure 2.**
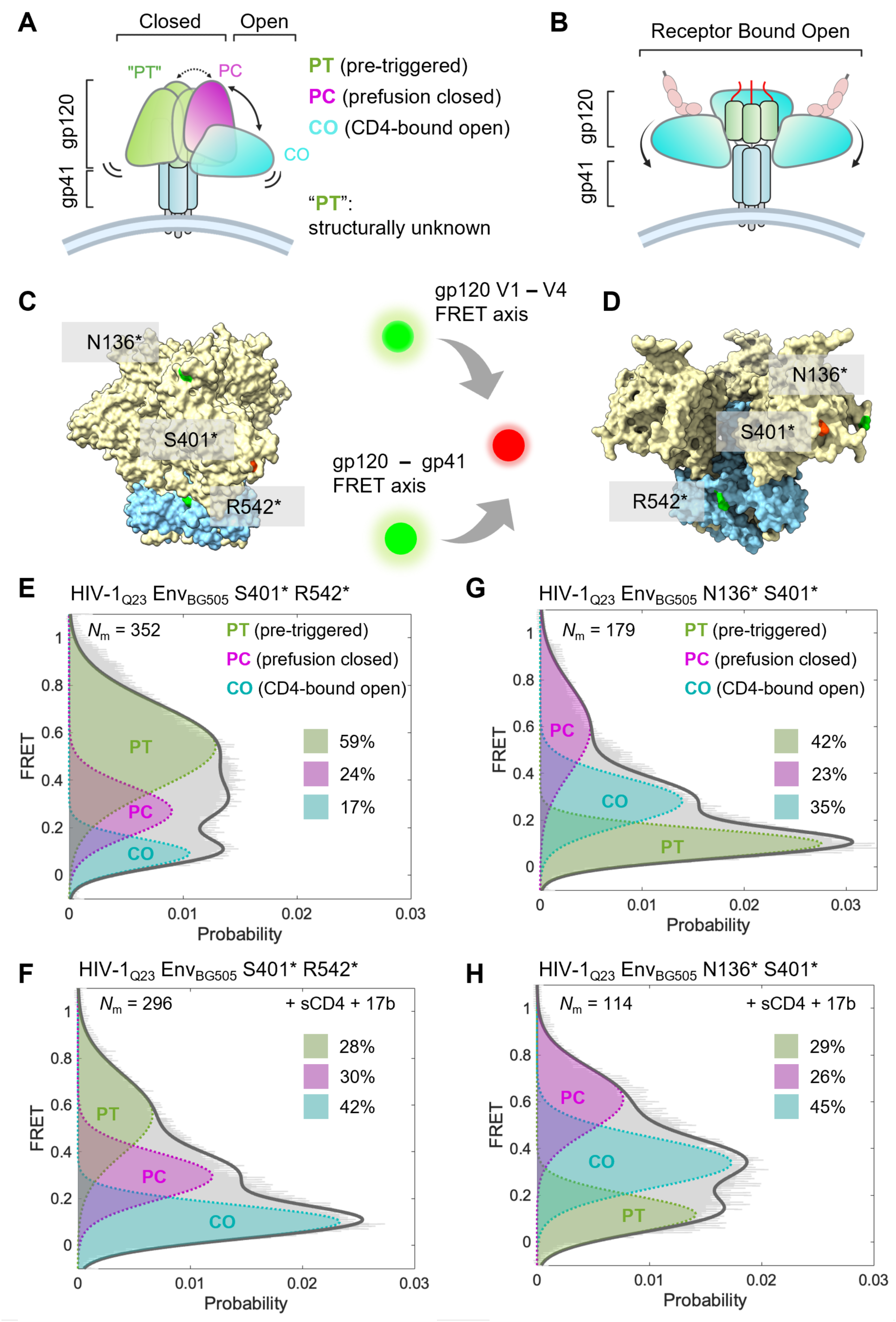
smFRET observation of global conformational opening of Env trimer from gp120–gp41 and gp120 V1V4 structural perspectives. **(A, B)** Schematic models of Env conformational states. (**A**) gp120 adopts three major conformations: “PT” (pre-triggered, structurally unknown), PC (prefusion closed), and CO (CD4-bound open), shown for a single protomer within the trimer, with the other two protomers remaining in the PT state. (**B**) Fully open Env trimer schematic showing all three gp120 subunits in the CD4-bound state. **(C, D)** Structural visualization of Env conformations from two smFRET probe perspectives. Env trimers in the prefusion closed (**C**, PDB #4TVP) and fully open (**D**, PDB #5VN3) states are shown with probe positions highlighted: gp120–gp41 (S401* R542*) and gp120 V1V4 (N136* S401*). **(E, F)** FRET histograms of Env_BG505_ S401* R542* (gp120–gp41 perspective) on native virions without ligand (**E**) or with soluble CD4 (sCD4) plus the co-receptor–mimicking antibody 17b (**F**). Ligand-free Env samples three primary conformational states (as in panel A), whereas sCD4 and 17b stabilize the fully open conformation (as in panel B). Here and elsewhere, *N_m_* indicates the number of FRET trajectories - molecules that were used to construct the histogram and were fit to the sum of three Gaussian distributions (see **Table S1**). State occupancies are reported as percentages. **(G, H)** Validation from the gp120 V1V4 perspective using Env_BG505_ N136* S401*. FRET histograms confirm a conformational shift toward the open-dominated population.

To quantify those populations observed from the S401–R542 axis, we compiled the overall conformational landscape and calculated state occupancies under ligand-free and ligand-bound conditions (**Fig. 2E–F**, **Table S1**). FRET histograms generated from hundreds of trajectories and fit with three Gaussian distributions revealed three well-separated populations, with high-FRET predominating in the absence of ligand. Addition of soluble CD4 (sCD4) and the co-receptor–mimicking antibody 17b shifted the distribution toward the low-FRET population (**Fig. 2F**, **S6**). Parallel smFRET measurements using the gp120 V1**–**V4 (N136*–S401*) probe pair (**Fig. 2G–H**) yielded similar ligand-free and ligand-induced shifts despite differences in absolute FRET values. These results collectively enabled the confident assignment of the low-FRET (~0.1), intermediate-FRET (~0.3), and high-FRET (~0.55) populations to the CO, PC, and PT states, respectively. As FRET values are inversely proportional to donor-acceptor separation, the two dyes are expected to be further apart in the CO state than in the PC state. This expectation is supported by atomistic molecular dynamics simulations, which show that Env adopts a larger inter-dye distance in the CO state than in the PC state (**Fig. S7**). Together, these results suggest the gp120–gp41 probe pair as a reliable reporter of Env conformational dynamics and CD4-associated opening. The three-state model consistently provided the simplest and most probable explanation for both our current data and previous studies.^17,24,27,47,66^ The consistency of these shifts across probe-pair positions provides compelling evidence that trimer opening is an intrinsic dynamic property of Env. It undergoes global conformational changes upon engaging with sCD4 and 17b, independent of the observed smFRET structural perspective.

To align our nomenclature with that of previous studies^17,19,24^, the PT state corresponds to previously designated State 1, the PC state corresponds to the symmetric State 2 (which the SOSIP-based soluble Env primarily adopts^66^), and the CO state corresponds to the fully open State 3. Additional conformational substates, including partially open or asymmetric Env trimers, or other low-abundance substates, may exist and may be resolved under different experimental or triggering conditions. For example, previous work showed that asymmetric intermediate conformations can be resolved using a heterotrimer design containing mixtures of wild-type and CD4-binding-deficient D368R protomers, where the State 2 FRET signal can originate from an unliganded protomer while the remaining one or two protomers are CD4-bound and open.^24^ In the present study, we did not use such a heterotrimeric design. Therefore, although our labeling strategy reports the major conformational populations of Env, it cannot unambiguously distinguish protomer asymmetric states, such as trimers with only one or two gp120 molecules open. Evidence from current and previous studies^17,19,24,47,66^ strongly supports the presence of three primary states of virus-associated Env, with additional substates that can be resolved under specific triggering conditions^23,27,67^. Accordingly, the three Gaussian models used here represent the three predominant conformational populations under our experimental conditions.

### Allosteric stabilization of Env in the prefusion closed state by gp120–gp41 interface and fusion peptide bNAbs

We next investigated how gp120–gp41 interface and FP bNAbs influence Env conformational dynamics. To this end, we performed smFRET analysis of fluorescently labeled Env_BG505_ S401*–R542* trimers on intact virions in the presence of 8ANC195,^68^ VRC34,^28,69^ or PGT151^58^ (**Fig. 3** and **Fig. S8**). Sequences of heavy and light chains of antibodies used in this study are provided in **Table S2**. Dose–response neutralization curves confirmed potent yet distinct activity of all three bNAbs against HIV-1_Q23_ Env_BG505_ virions (**Fig. 3A**, left). The three antibodies engage distinct epitopes at the gp120–gp41 subunit interface (**Fig. 3A**, right)^28,58,68,69^: 8ANC195 and PGT151 make extensive contacts with both gp41 and gp120, whereas VRC34 binds predominantly to the N-terminal region of the fusion peptide. These distinct binding modes likely result in differential effects on the conformational landscape of native Env, motivating smFRET studies to uncover their impact on Env dynamics.

**Figure 3.**
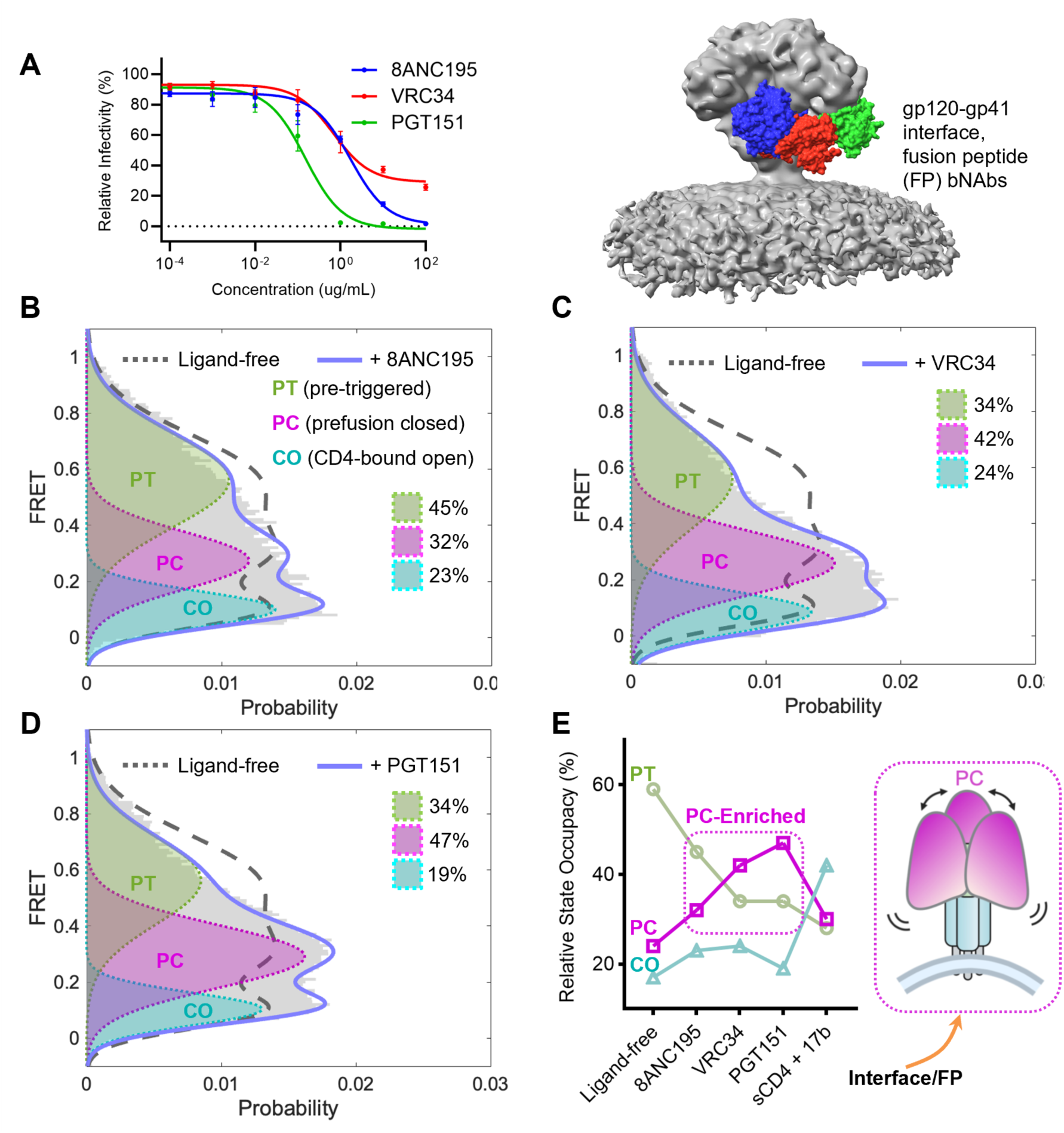
Stabilization of HIV-1 Env in the prefusion closed state by gp120–gp41 interface or FP bNAbs. (**A**) Dose–response neutralization curves (left) of HIV-1_Q23_ Env_BG505_ by the gp120–gp41 interface or FP bNAbs 8ANC195, VRC34, and PGT151, with binding epitopes mapped onto the membrane-bound Env trimer (EMD-21412, right). (**B**) FRET histogram of HIV-1_Q23_ Env_BG505_ S401* R542* in the presence of 8ANC195, overlaid with ligand-free Env (dashed gray), showing modest conformational shifts toward downstream states. (**C**, **D**) FRET histograms for VRC34 (**C**) and PGT151 (**D**), as in **B**, revealing redistribution of Env populations from pre-triggered (PT) to prefusion closed (PC) state dominance. (**E**) Line graph showing relative state occupancy of Env under different antibody conditions, demonstrating progressive enrichment of the PC state by 8ANC195, VRC34, and PGT151.

We compiled FRET histograms (**Fig. 3B–D**), which revealed antibody-specific effects on Env conformational sampling. Histograms were constructed from ***N***_m_ of smFRET trajectories, as indicated in each plot. Representative examples (**Fig. S8A–C**) displayed characteristic anti-correlated donor–acceptor intensity fluctuations and state-to-state transitions, consistent with conformational sampling between PT, PC, and CO states. In the presence of 8ANC195, Env displayed a modest redistribution from the high-FRET PT state (occupancy decreased from 59% to 45%) toward the intermediate-FRET PC (24% to 32%) and low-FRET CO (17% to 23%) states. By contrast, VRC34 and PGT151 induced more pronounced shifts, resulting in PC state dominance (42% and 47%, respectively) and a marked reduction in PT occupancy (54% to 34%). We further summarized state occupancies under antibody-incubated conditions, in reference to those for ligand-free and CD4/17b-incubated conditions, using bar (**Fig. 3E**, left) and line graphs (right) derived from quantitative model fitting of FRET histograms with constrained three-state Gaussian models (**Table S1**). These analyses revealed allosteric enrichment of the PC state across the antibody panel (**Fig. 3E**, right inset model plot), with the strongest stabilization observed for PGT151 (47%). Ligand-induced stabilization, as revealed by smFRET here, reflects a redistribution of the conformational ensemble while preserving the intrinsic dynamic nature of Env. These results suggest that gp120–gp41 interface and FP-targeting bNAbs remodel the Env conformational landscape by allosterically shifting the distribution toward the prefusion-closed state along the gp120 opening pathway.

### MPER-directed bNAbs allosterically stabilize the prefusion-closed state, while 10E8.4/iMab exerts a dual effect by also promoting the CD4-bound open state

We next examined how MPER-directed bNAbs DH511.2_K3,^70^ VRC42,^71^ and the bispecific antibody 10E8.4/iMab influence Env conformational dynamics on native virions. Sequences are listed in **Table S2**. All antibodies potently neutralized HIV-1_Q23_ Env_BG505_, with the bispecific engineered 10E8.4/iMab,^38,39^ which combines a variant of the 10E8 antibody with the CD4 receptor-targeting antibody iMab, exhibiting exquisite potency (**Fig. 4A**, left). Their distinct binding epitopes, located distal to the gp120 opening apex, were defined from Env–bNAb complex structures solved with truncated gp41 peptide constructs and mapped to the MPER region of Env (**Fig. 4A**, right), which lies adjacent to the viral membrane. DH511.2_K3 is a 10E8-like antibody,^70^ and VRC42 shares similar binding epitopes with the 4E10 antibody.^71^ For 10E8.4/iMab, neither the structural details of its complex with Env nor its impact on Env dynamics has been defined, although the structure of 10E8 itself with MPER peptide has been determined.^72^ In light of this, we used smFRET to directly observe, from the gp120-gp41 axis (S401* R542*), how these antibodies allosterically remodel Env conformational distributions in the native viral context.

**Figure 4.**
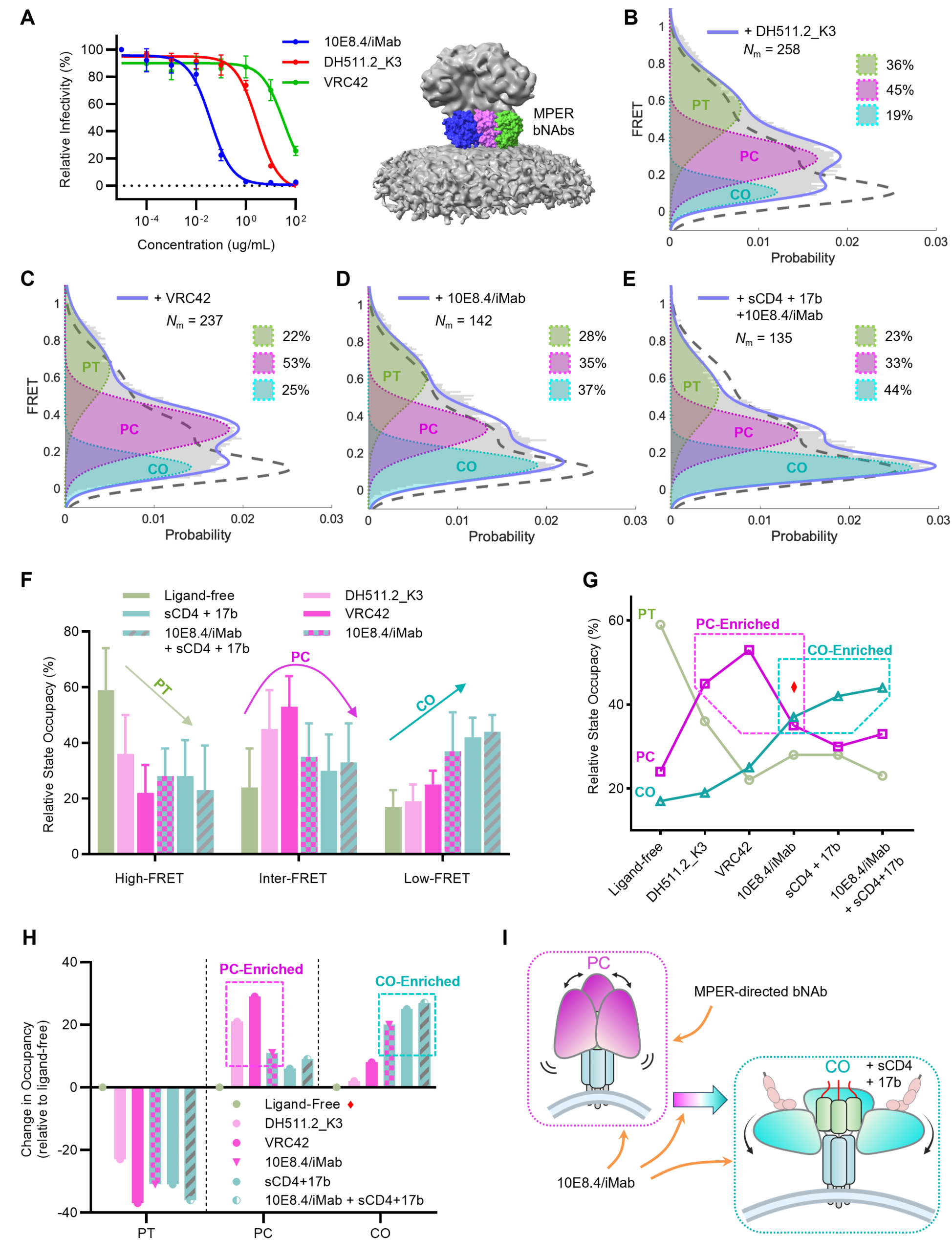
MPER-directed bNAbs enrich Env in the prefusion-closed (PC) state, while 10E8.4/iMab also enriches Env in the CD4-bound open (CO) state. (**A**) HIV-1 neutralization by MPER-directed antibodies DH511.2_K3 and VRC42, and 10E8.4/iMab. Left: neutralization curves; right: MPER binding epitopes on Env. (**B, C**) FRET histograms of Env_BG505_ S401* R542* in the presence of DH511.2_K3 (**B**), or VRC42 (**C**). These MPER-directed bNAbs shift Env toward dominance of the PC state. The FRET histogram for Env with sCD4 and 17b (dashed gray) was included in each plot for reference. **(D, E**) smFRET results for bi-valent 10E8.4/iMab alone (**D**), which shifts Env further downstream along the opening pathway with a broad range across PC and CO, and with sCD4 and 17b (**E**), which further stabilizes Env in the open CO state. (**F, G**) Bar graph (**F**, mean ± s.e.m.) and line graph (**G**) showing relative state occupancy of Env under different antibody-binding conditions, demonstrating shifts from PT to PC with MPER bNAbs, and progression toward CO with 10E8.4/iMab alone or combined with sCD4 and 17b. (**H**) Bar graph of changes in state occupancy of Env exerted by MPER antibodies relative to ligand-free. (**I**) A derived schematic summarizing smFRET results of preferential conformations of Env targeted by MPER-directed bNAbs and 10E8.4/iMab.

Two MPER-directed bNAbs exhibited a preference for the intermediate-FRET state, and thus a prevalent residence in the PC state observed from FRET histograms (**Fig. 4B–C**). The conformational redistribution from PT to PC dominance for DH511.2_K3 (**Fig. 4B**) and VRC42 (**Fig. 4C**) suggests that MPER engagement allosterically restricts the trimer apex from fully opening by stabilizing the base where the binding epitopes reside. Otherwise, one would expect an increase in the CO state rather than no notable change. The result is somewhat unexpected. We initially hypothesized that Env would need to adopt a more open conformation to permit bNAb access to the sterically restricted epitopes near the viral membrane. The results, nevertheless, are consistent with those obtained from cryo-EM structures using the Env ectodomain or the isolated MPER domain.^70,71^ There are three plausible mechanisms by which MPER-directed antibodies could access their epitopes: (1) partial or full gp120 opening to relieve steric constraints on gp41, (2) trimer tilting relative to the viral membrane, and (3) local membrane bending or curvature. Although our initial hypothesis favored some degree of gp120 opening, the observed stabilization of the PC state suggests that MPER accessibility may arise through trimer tilting or membrane deformation, in line with a recent observation of membrane-coupled Env trimer tilting on virions.^73^

The bivalent 10E8.4/iMab, strikingly, induced a distinct conformational response of Env on virions. It stabilized Env in both PC and CO states, with the latter predominating at the cost of the PT state (**Fig. 4D**). FRET histograms revealed a redistribution beyond PC toward the largest increase in CO-state sampling (**Fig. 4D**), When combined with sCD4 and 17b, 10E8.4/iMab further amplified this shift, driving Env even further into the CD4-bound open state with reduced sampling of other conformations (**Fig. 4E**). Therefore, unlike the other two monovalent MPER antibodies that primarily stabilize the PC state, the results indicate that 10E8.4/iMab exerts a dual effect (**Fig. 4F*–*I**), favoring both the PC and, more profoundly, the CO states at the expense of the PT state. The order of antibody presentation in quantitative results and comparisons of state occupancies (**Fig. 4F–H**) was intentionally arranged to illustrate this continuum of Env opening, highlighting the gradual redistribution of state occupancies from MPER-bNAb–stabilized closed conformations to 10E8.4/iMab- and sCD4/17b-driven open states.

We next performed smFRET of Env in the presence of 10E8.4 and iMab individually to examine their separate conformational effect on the virus-associated Env (**Fig. S9**). We found that 10E8.4 behaves similarly to other MPER-directed antibodies in enriching the PC state, whereas iMab alone does not appear to have a notable effect on the conformational propensity of Env (**Fig. S9**).

The distinct conformational responses induced by 10E8.4/iMab are intriguing and unexpected, given that the 10E8 arm behaves similarly to other MPER-directed antibodies, and the iMab arm engages CD4 on the host membrane (and thus is not expected to interact with Env trimer). Notably, this effect of favoring the CO state was observed even in the absence of CD4 or the host membrane context. The unique neutralization of 10E8.4/iMab is reminiscent of CD4-induced opening of the spike, and indeed, the binding of MPER-directed antibodies has been shown to synergize with that of CD4,^74,75^ as explored further in the Discussion section.

### Antibodies shift Env transition frequencies with limited effects on rates while preserving the opening pathway

To investigate how antibodies influence Env dynamics beyond overall conformational distributions and steady-state occupancies, we analyzed single-molecule trajectories to extract state-to-state transitions and quantify their kinetics (**Methods**). Transition density plots (TDPs) were generated by plotting the initial versus final FRET values for every detected transition, with color intensity reflecting the relative frequency of transitions (transitions per second) (**Fig. 5A**). These plots report which transitions occur and how often, with the total number of transitions (*N*_t_) and molecules analyzed (*N*_m_, the same molecules used for the FRET histograms and occupancy analysis) indicated on each plot. These TDPs revealed antibody-dependent redistribution of state-to-state transitions: interface/FP bNAbs (such as PGT151) and MPER bNAbs (such as DH511.2_K3) enriched transitions into and within the intermediate-FRET PC state, whereas 10E8.4/iMab and sCD4/17b increased transitions leading toward the low-FRET CO state from both PT and PC. The expected set of allowable transitions between PT and PC, between PC and CO, and rarely between PT and CO was preserved under all conditions (**Fig. 5A**), suggesting that the overall order and directionality of Env state-to-state transitions remain unchanged.

**Figure 5.**
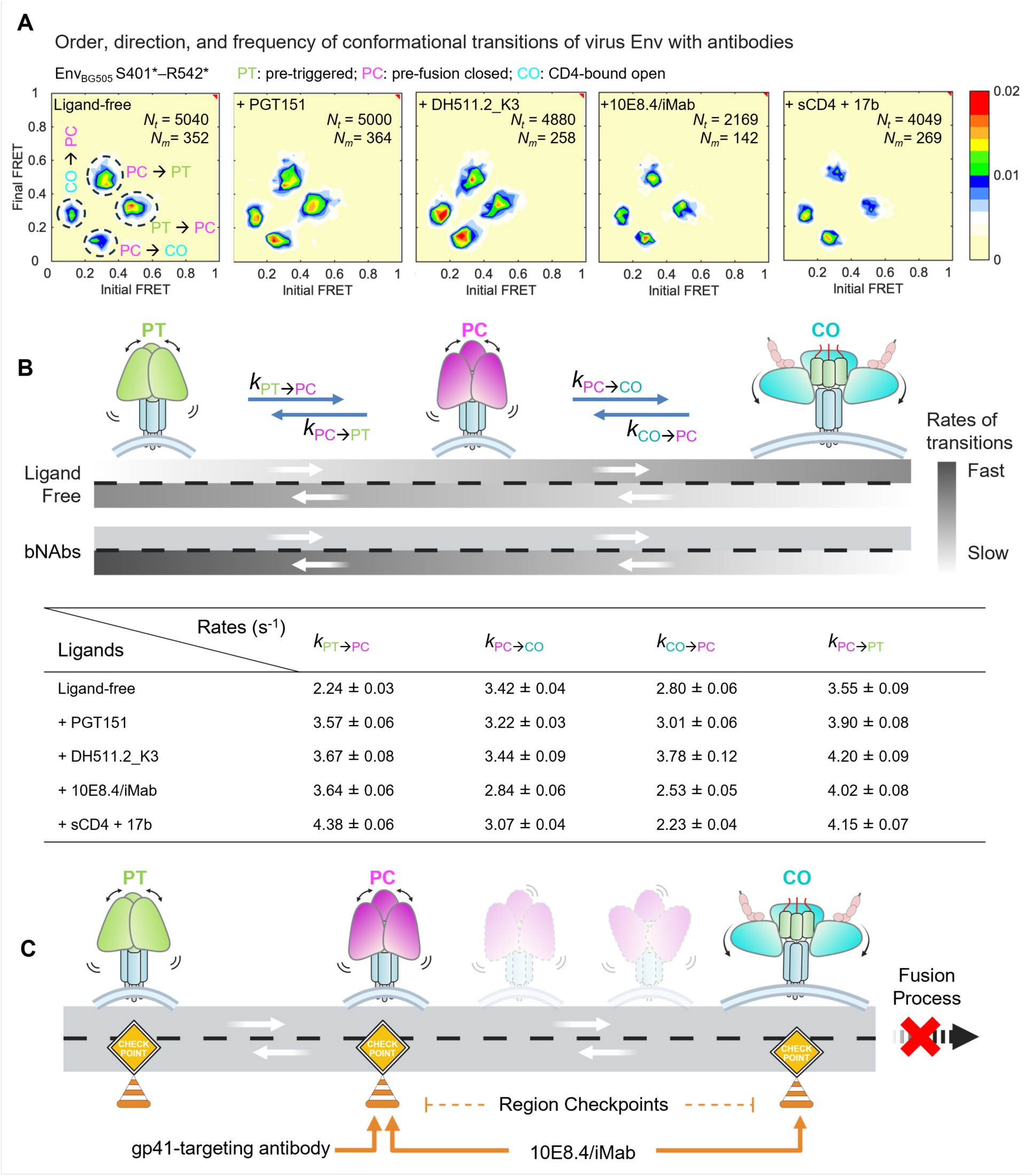
Kinetic analysis of Env transitions exerted by gp41-directed antibody and 10E8.4/iMab, and an integrative model depicting allosteric control across conformational states. (**A, B**) gp41-directed antibody and the bispecific 10E8.4/iMab alter Env transition frequencies with slight effects on kinetic rates while preserving the opening pathway. (A) Transition density plots (TDPs) showing the order, directionality, and frequency of Env conformational transitions in the presence of representative bNAbs. Each TDP plots the initial versus final FRET value for every detected transition, with color intensity indicating the relative frequency of those transitions (transitions per second). *N*_t_ = total number of transitions from *N*_m_ (total number of molecules), indicated on each plot. These plots reveal antibody-dependent redistribution of state sampling among PT, PC, and CO conformations. (B) Schematic (top) and table (bottom) summarizing transition rates between allowable Env conformations in the presence of interface/FP, MPER, and 10E8.4/iMab antibodies. (**C**) Integrative working model of gp41 antibody-mediated allosteric control of Env conformational states. Native Env on virions transitions through three primary states (PT, PC, and CO) along the opening pathway toward fusion, depicted as sequential checkpoints along the opening/fusion road targeted by antibodies. Most gp41-directed bNAbs act preferentially at the PC checkpoint, whereas 10E8.4/iMab can act at both PC and CO. The MPER- and CD4-binding arms of 10E8.4/iMab may function as a mechanical brace, restraining Env from breaking free at PC, CO, and the secondary checkpoints between them, which could explain its superior potency.

To determine whether these occupancy shifts were accompanied by changes in transition kinetics, we constructed survival probability plots for each FRET-defined state (**Fig. S10**). Dwell times for molecules in each state prior to a transition were pooled, and exponential functions were fit to the resulting distributions to derive weighted-average rates for transitions between allowable conformations. These rate constants, which fall on the scale of seconds, were summarized as a directional connectivity schematic and a numerical table (**Fig. 5B**). This analysis revealed only modest antibody-induced changes in transition rates, suggesting that the primary effect of the tested antibodies is to allosterically redistribute the frequencies of state sampling rather than fundamentally rewire the intrinsic kinetics of Env motion. Together with the TDPs, these results demonstrate that antibodies shift transition frequencies while preserving the overall network of allowable transitions, leaving the canonical opening pathway intact.

## Discussion

### Two modes of neutralization via allosteric control of Env dynamics across distinct conformational checkpoints

In this study, we demonstrate that gp41-directed bNAbs exert long-range allosteric control of Env dynamics through two different conformational outcomes. Most bNAbs targeting the gp120–gp41 interface, FP, and MPER stabilize the prefusion-closed (PC) state and restricted sampling of the pre-triggered (PT) conformation. In contrast, the bispecific 10E8.4/iMab, a promising therapeutic currently in clinical trials, exerts a dual effect by also stabilizing the CD4-bound open (CO) state. These findings lead to our current working model (**Fig. 5C**). The native Env on virions transitions through three primary conformational checkpoints, from PT through PC to CO, along the opening pathway towards fusion. Multiple secondary checkpoints likely exist between the PC and CO states. Whereas most gp41-directed bNAbs act primarily at the PC checkpoint, 10E8.4/iMab can engage both PC and CO checkpoints and may also act at substates in between.

These conformational controls over Env reveal two dynamic modes of neutralization: (**1**) allosteric stabilization of the prefusion-closed state, and (**2**) premature triggering into the open state. Regarding the second mode, associated with 10E8.4/iMab, we cannot exclude the possibility that the CD4-targeting arm enhances or stabilizes binding of the 10E8.4 arm to Env, thereby improving access to its epitope in the CD4-bound conformation. Premature triggering of global conformational changes and localized epitope binding are not mutually exclusive; rather, they may coexist. In either case, both modes reflect the intrinsic allosteric regulation of Env, where perturbations at one site, such as antibody binding or a single amino acid substitution, can propagate across the trimer to stabilize or destabilize distant domains. Prior studies have shown that even a single or a few substitutions can reconfigure distal structural elements,^19,20,22,76,77^ underscoring the remarkable sensitivity of Env to allosteric perturbation.

These two modes form a continuum, with different antibodies shifting Env among checkpoints along the opening pathway (**Fig. 5C**). Such dynamic control likely underlies the exceptional potency of some antibodies, such as the bispecific 10E8.4/iMab, by broadening the range of neutralized conformations while reducing opportunities for viral escape across multiple conformational checkpoints.

### Striking Env opening by 10E8.4/iMab in the absence of CD4

The finding that MPER-directed bNAbs stabilize the PC state also suggests that MPER accessibility on gp41 does not necessarily require trimer opening, which was unexpected, as discussed in the Results section. Even more surprising, however, was the observation that 10E8.4/iMab stabilizes the CO state in addition to the PC state, despite the absence of soluble or membrane-associated CD4.

We suspect that the two binding arms of 10E8.4/iMab may act like a molecular (mechanical) brace on a hinge, limiting Env escape at and between these two primary conformational checkpoints (**Fig. 5C**). This effect likely broadens the range of conformational states and substates accessible to neutralization, which could explain its superior neutralizing abilities in both potency and breadth. Whether 10E8.4/iMab can target multiple unknown substates or secondary checkpoints (**Fig. 5C**) of Env along the opening trajectory remains to be clarified; however, two recent integrative studies provide relevant context. Using collective molecular dynamics, one computational study revealed a previously overlooked neutralization-relevant occluded-intermediate state between PC and CO that emerged in the presence of one of four specific bNAbs.^27^ This newly identified state was further supported by smFRET analyses, which showed reproducible conformation-related shifts validated through statistical evaluation of multiple models.^27^ Another recent cryo-EM study identified several open intermediates with varying degrees of fusion peptide accessibility, induced by bNAb VRC34 under different open-triggering conditions.^28^ Again, the existence of these new intermediates was supported mechanistically by smFRET results using new probes performed under the same conditions.^28^ Of note, these newly identified conformations or states between PC and CO were not previously observed in initial cryo-EM^69,78^ or smFRET^69^ of Env–VRC34. These observations are encouraging, as they underscore the likelihood that 10E8.4/iMab targets a broad range of Env conformations, including PC, CO, and even currently uncharacterized “hidden” Env conformations. Future structural investigations, particularly high-resolution studies of 10E8.4/iMab bound to membrane-associated Env and membrane-embedded CD4, could provide deeper mechanistic insights into how distinct binding modes mediate allosteric regulation and inhibit fusion through conformational control, trimer tilting, or membrane deformation.

### Underappreciated transition kinetics of Env dynamics

An additional mechanistic insight, often underappreciated, is that antibody-induced conformational redistribution occurred without major rewiring of the intrinsic opening pathway (**Fig. 5A–B**). Our kinetic analysis reveals that the overall transition network and directionality (PT ↔ PC ↔ CO), which describe the progression from pre-triggered to prefusion closed and onward to CD4-bound open, remain preserved. Antibody effects were primarily reflected in shifts in transition frequencies and state occupancies, with only modest changes in transition rates. This suggests that antibodies primarily act by stabilizing or destabilizing specific states within the preexisting pathway, rather than creating alternative routes. Such long-range allosteric modulation provides a mechanism for neutralization while maintaining the intrinsic connectivity of Env dynamics.

### Coherent mechanistic landscape linking Env structures and dynamics to neutralization

The observed conformational redistributions of Env by 8ANC195, VRC34, PGT151, DH511.2_K3, VRC42, and 10E8.4/iMab are mechanistically consistent with structural studies that mapped Env–antibody epitopes on predominantly closed conformations.^28,58,68–72,79^ They are also in line with previous smFRET studies using probes confined to gp120,^28,47,66,69^ as well as molecular virology studies examining neutralization sensitivity or resistance in tier-level viruses or those with Env modifications.^37,74,75,80,81^ Structural analyses have revealed how Env–antibody complexes neutralize through direct binding, identifying atomic footprints and steric blockades that prevent CD4 or coreceptor engagement or insert into fusion-critical elements to block entry.^28,58,68–72,79^ These static structural snapshots of bNAb–Env complexes, primarily captured with engineered or truncated Env in the closed conformation, align with the PC dominance observed in this study. Previous smFRET analyses of PGT151 and VRC34 with probes restricted to gp120^28,47,66,69^ revealed their preferential recognition of PC, distinct from the pre-triggered (PT) state. Our updated smFRET approach, using dual bioorthogonal click labeling across the gp120–gp41 structural axis, reinforces the finding of PC stabilization and provides additional insight into transition kinetics. The smFRET characterization of 8ANC195, VRC42, DH511.2_K3, and 10E8.4/iMab presented here is, to our knowledge, the first of its kind, revealing Env conformational responses and transition kinetics. Molecular virology studies have also reported correlations between the neutralization sensitivity of gp120- and gp41-directed bNAbs and the conformational flexibility, and, in some cases, the degree of opening or closing of Env trimers.^37,74,75,80,81^ Our smFRET data, which capture Env conformational distributions and bNAb-induced shifts, reveal intrinsic conformational flexibility consistent with these studies in principle. In particular, one MPER-directed bNAb, 10E8.4/iMab, can open the spike in a manner similar to CD4, consistent with other experiments showing that MPER binding and CD4 binding synergize.^74,75^

Overall, our results introduce dynamic and kinetic dimensions to the neutralization landscape within the native viral context. Our data reveal the dynamic range of Env responses to antibody binding, highlighting the complementary aspects of structural stabilization and conformational plasticity in neutralization. These findings provide new mechanistic insights into neutralization by identifying how antibodies target Env on virions across multiple conformational checkpoints through mediating its conformational sampling probabilities, transition frequencies, and the trajectories it undergoes.

### Complementary views from smFRET and structural studies in general

The smFRET analyses reveal that bNAb binding generally shifts the conformational equilibrium of Env rather than driving all trimers into a single structural state. Even in the presence of saturating ligands, substantial fractions of Env remain in alternative conformations, highlighting the intrinsic conformational heterogeneity of the native viral spike. This heterogeneity likely reflects both differences in ligand binding and the dynamic energy landscape of Env. In this regard, smFRET and structural methods provide complementary information. While cryoEM and cryoET define the molecular architecture of individual conformational states, smFRET quantifies their relative populations and dynamic interconversion. The conformational ensembles observed by cryoEM may also be influenced by factors such as conformation-dependent radiation sensitivity and the requirements of image classification, making low-abundance or highly dynamic states more difficult to recover. Together, these complementary approaches provide a more complete view of Env conformational dynamics than either method alone. The transition density plots further illustrate how different ligands reshape the conformational landscape by altering both state occupancies and the transitions between them, rather than simply stabilizing a single static structure.

### Limitations of this study

Several limitations should be acknowledged. Our smFRET studies were performed on virion-associated full-length Env, but in the absence of host membranes. This limits interpretation, as host lipid composition, membrane curvature, and host factors could influence Env dynamics and potentially contribute to neutralization. We did not capture the full conformational continuum, particularly downstream states after coreceptor engagement and gp41 refolding, which will require dual labeling of gp41 and host membranes together with complementary methods such as time-resolved cryo-EM or cryo-ET. Photobleaching of the donor and acceptor fluorophores within approximately 10 seconds of excitation limits the observation window for individual molecules. Technically, while smFRET provides millisecond-scale resolution of state sampling suitable for Env dynamics, very short-lived conformations remain underrepresented. Labeling at the gp120–gp41 axis was optimized to minimize perturbation, but chemical modifications could still influence local structural fluctuations normally below our detection threshold. The conformational effects of antibodies were examined in a single Env background (BG505, widely used and representative), although Env sequence diversity may modulate antibody-dependent effects. Finally, the link between Env dynamics and neutralization potency will require integration with in-depth structure–function analyses in a native context. Nevertheless, our findings refine the mechanistic understanding of antibody-mediated neutralization and reveal gp41 as a promising target for strategies that exploit the intrinsic allosteric vulnerabilities of Env.

## Materials and methods

### Cell lines and cell maintenance

HEK293T cells (ATCC # CRL-3216) were used for producing replication-defective HIV-1 viruses. TZM-bl cells (BEI Resources # HRP-8129), which stably express the receptor CD4 and the co-receptor CCR5, were used as target cells to quantify the infectivity of the produced HIV-1 viruses and to assess the neutralizing activity of antibodies. These cell lines were cultured in high glucose Dulbecco’s Modified Eagle Medium (Gibco # 11965-092) supplemented with 10% (v/v) heat-inactivated fetal bovine serum (Gemini Bio # 100-106), 100 mg/mL penicillin-streptomycin (Gibco # 15140-122), and 6 mM L-glutamine (Gibco # 25030-081), and maintained in a 37°C incubator supplied with 5% CO_2_.

### Plasmid construction

Tagged Env plasmids used in this study were constructed by site-directed mutagenesis using the Amber-free HIV-1_Q23_ Env_BG505_ ΔRT (Addgene # 213007)^47^ plasmid as a template. The plasmid HIV-1_Q23_ Env_BG505_ carries the gene encoding the Env protein from the BG505 strain, while all non-Env HIV-1 genes are derived from the HIV-1 Q23 strain.^82,83^ Plasmid HIV-1_Q23_ Env_BG505_ ΔRT was made by deleting the gene encoding the reverse transcriptase (RT) from the parental construct.^24^ The amber-free HIV-1_Q23_ Env_BG505_ ΔRT (Addgene # 213007) plasmid was created from HIV-1_Q23_ Env_BG505_ ΔRT by substituting all TAG (amber) stop codons in the Pol, Vif, Vpu, and Rev genes with TAA (ochre) stop codons.^47^ The plasmid amber-free HIV-1_Q23_ Env_BG505_ S401^TAG^ R542^TAG^ was generated by introducing TAG stop codons at the codons corresponding to Ser401 and Arg542 within the Env sequence. The dual-amber Env_BG505_ S401^TAG^ R542^TAG^ construct was generated using a similar mutagenesis approach. All plasmids were amplified in Stbl3 competent cells (Invitrogen, #C7373-03) and verified by DNA sequencing prior to use.

### Virus production

The preparation of HIV-1_Q23_ Env_BG505_ viral particles has been previously described.^24,47,66^ Briefly, healthy, exponentially growing, and mycoplasma-free HEK293T cells were used to produce viral particles. One day prior to transfection, HEK293T cells were seeded into culture plates to ensure that cell confluency exceeded 70% at the time of transfection. The growth medium was then replaced with Opti-MEM, and the transfection complex was prepared by mixing polyethylenimine (PEI; 1 mg/mL) with plasmid DNA (amber-free HIV-1_Q23_ Env_BG505_ or its derivatives) at a 3:1 (v:w) ratio of PEI to total plasmid. The mixture was incubated at room temperature for 15 minutes before being added to the cells. For packaging HIV-1 viral particles carrying dual-amber mutant Env (such as N136^TAG^ S401^TAG^ or S401^TAG^ R542^TAG^),^47^ an additional plasmid, tRNA^pyl^/NESPyIRS^AF^ (a gift from the Edward Lemke Lab),^41,50^ was co-transfected to enable incorporation of the noncanonical/unnatural amino acid (ncAA). This plasmid encodes an aminoacyl-tRNA synthetase (aaRS) that incorporates ncAA into its cognate tRNA and was included at a 3:1 ratio relative to the Env plasmid. The ncAA trans-cyclooct-2-en-L-lysine (TCO*A; SiChem #SC8008) was added to the culture medium at a final concentration of 250 μM. For the preparation of viral particles used in subsequent infectivity and neutralization assays, an additional plasmid, HIV-1-inGluc, encoding the secreted Gaussia luciferase (Gluc) enzyme, was included in the transfection mixture. Four to six hours after transfection, the transfection medium was replaced with complete growth medium, and the cells were incubated for 40 hours. The culture supernatant containing viral particles was collected, filtered through a 0.45 μm membrane filter (PALL #4654), and concentrated by ultracentrifugation through a 15% (w:v) sucrose cushion in PBS at 25,000 rpm for 2 hours using an SW28 rotor (Beckman Coulter). The resulting viral pellets were resuspended in PBS for subsequent use.

### Western blotting

Virus particle preparations or cell lysates were mixed with Laemmli SDS sample buffer (Thermo Scientific, #J61337-AD) and heated at 95 °C for 15 minutes. The denatured protein samples were separated on a 4–12% Bis-Tris gradient gel by SDS-PAGE and subsequently transferred onto a PVDF membrane using the Trans-Blot Turbo system (Bio-Rad, #1704150). The membrane was blocked with 5% (w:v) non-fat milk (RPI, #M17200) in TBST buffer at room temperature for 1 hour, followed by incubation with the primary antibody against HIV-1 gp120 (BEI Resources, #ARP-288) diluted in 5% non-fat milk/TBST at 4 °C overnight. After washing the membrane three times with TBST (10 minutes each), it was incubated with HRP-conjugated anti-sheep IgG secondary antibody (Proteintech, #SA00001-16) at room temperature for 1 hour. The membrane was washed again three times with TBST, and protein bands were visualized using a chemiluminescent HRP substrate (Millipore, #WBKLS0500). Signal detection and quantification were performed with the ChemiDoc Imaging System (Bio-Rad) using Image Lab software.

### Production of soluble CD4 and antibodies

The following monoclonal antibodies and proteins were expressed in Expi293 cells by transient transfection of heavy-light chain plasmids, followed by protein A affinity column, size exclusion chromatography, and buffer exchange to 20 mM PBS pH 7.5, 0.002% w/v azide, and then flash-frozen for use. Overall, antibodies 17b, DH51.2_K3, 8ANC195, VRC34, VRC42, PGT151, 10E8.4, iMab, and 10E8.4/iMab were produced in the same way. sCD4 was also expressed transiently and purified as described previously.^28^

### HIV-1 infectivity and antibody neutralization

Infectivity and neutralization assays were performed as previously described^47^. One day before the assay, TZM-bl cells were seeded into 96-well culture plates. For virus neutralization assays, antibodies were serially diluted in basic DMEM and transferred to a new 96-well plate at 100 μL per well, with four replicates for each concentration. Subsequently, 50 μL of virus solution was added to each well, mixed thoroughly, and incubated at 37 °C for 1 hour. For infectivity assays, no antibody was added, and the virus solution was applied directly to the cells. After incubation, the culture medium from the TZM-bl cells was removed and replaced with 50 μL per well of DMEM containing 20% FBS. The antibody–virus mixtures (or virus alone for infectivity assays) were then added to the cells and incubated for 48 hours at 37 °C. Following incubation, the plates were gently mixed, and 50 μL of the supernatant from each well was transferred to a white microtiter plate. Luminescence was measured using a BioTek Synergy H1 microplate reader equipped with an automatic injector, following the manufacturer’s instructions for the Gaussia Luciferase Glow Assay Kit (Thermo, #16161). Data analysis was performed using GraphPad Prism software.

### Diameter measurement of HIV-1 viral particles

The diameter distribution of HIV-1 virions diluted in BBS buffer was measured by nanoparticle tracking analysis (NTA) using a ZetaView X30 instrument (Particle Metrix). After sample loading, the instrument automatically recorded videos across three cycles and 11 positions to track particle motion and calculate their hydrodynamic diameter and concentration based on the Stokes–Einstein equation. Data acquisition and analysis were performed using ZetaView software (version 8.05.16 SP3).

### Cryo-ET sample preparation and tomogram reconstruction

Virus particles for the Cryo-ET imaging were produced using the plasmid carrying the RT gene. After the supernatant containing HIV-1 virions was collected and filtered, the aldrithiol-2 (AT-2, Sigma # 143049) was added at a final concentration of 0.5 M and incubated at 4°C with rotation for 12-18 hours to inactivate the HIV-1 virus. The viral particles are then purified by sucrose density gradient centrifugation. Viral particles were mixed with a 6 nm gold tracer (Aurion). 4 μL of the mixture was placed onto freshly glow-discharged 200 mesh Cu Quantifoil R 2/1 grids, blotted for 5 s and plunge frozen in liquid ethane by using Vitrobot Mark IV (FEI Co.). Vitrobot was maintained at 4°C and 100% humidity during all these experiments. Frozen grids were imaged using a 200 kV Glacios Selectris X with a Falcon 4i direct electron detector. Tilt-series were collected using a dose-symmetric tilting scheme from −51° to +51° with a step size of 3°, and Tomography 5 software (Thermo Fisher Scientific) was employed at approximately 5 μm defocus. Tilt-series were collected at a magnification of 63000X, corresponding to a pixel size of 2.01 Å per pixel. The total dose per tilt series was ~80e^−^/Å^2^ distributed over 35 stacks. Each stack contains approximately ten images. We used IMOD to facilitate data processing, which includes drift correction of dose fractionated data and assembly of corrected sums into tilt series, automatic fiducial seed model generation, alignment, and contrast transfer function correction of tilt series^84^ and weighted back projection (WBP) reconstruction of tilt series into tomograms using Tomo3.^85^

### Preparation and fluorescent labeling of virus Env for smFRET

Viral particles used for smFRET analysis were RT-deleted and prepared using a method similar to that described previously.^47^ S401* R542* virions refer to dual-color, fluorescently labeled particles generated by incorporating noncanonical amino acids (ncAAs) at amber (TAG) codons corresponding to S401 on gp120 and R542 on gp41 via amber suppression, followed by fluorophore conjugation through click chemistry. N136* R542* virions were generated analogously.

During transfection, plasmids encoding amber-free HIV-1_Q23_ Env_BG505_ ΔRT and those carrying dual-amber Env variants (S401^TAG^ R542^TAG^ or N136^TAG^ S401^TAG^) were mixed at a calculated ratio (4:1) based on their relative amber suppression efficiencies (~20%) of S401^ncAA^ R542^ncAA^ or N136^ncAA^ S401^ncAA^ Env on virions. By assuming random assembly of Env trimers, the use of a very diluted Env ratio (untagged Env: tagged Env = ~20:1) will ensure that most virions carrying wild-type Env trimers, and among those virions carrying a single dual-ncAA gp120 within an otherwise wild-type Env background could be subsequently fluorescently labeled and identified using single-molecule imaging.

Virus pellets were resuspended in labeling buffer containing 50 mM HEPES, 10 mM MgCl₂, and 10 mM CaCl₂. Fluorescent labeling of Env by click chemistry was performed as previously described.^42,47^ Briefly, amber-free virions containing click-reactive TCO*A residues were incubated in a reaction mixture containing 0.1 mM tetrazine-conjugated Cy3 derivative (LD555-TTZ, Lumidyne) and Cy5 derivative (LD655-TTZ, Lumidyne) fluorophores at room temperature for approximately 6 hours. The reaction was quenched by the addition of 1 mM BCN-OH followed by incubation for 10 minutes. Subsequently, PEG2000-biotin was added to a final concentration of 0.1 mg/mL, and the mixture was incubated for 30 minutes at room temperature with rotation. Excess dye and lipids were removed by ultracentrifugation through a 6–18% Optiprep gradient at 40,000 rpm for 1 hour. Of note, clickable dyes can be randomly coupled to two ncAA TCO*A in Env trimer on the virus. Unlabeled and multi-labeled virions can be easily identified based on the correlation between synchronized two-channel fluorescence signals (as described below). The fluorescently labeled virions were then stored at –80 °C until use.

### Excitation and emission spectra characterization of virions

Viral particles were produced from the dual-amber Env S401^TAG^ R542^TAG^ construct using the methods described above. The concentrated viral preparation was divided into three equal aliquots and subjected to fluorescent labeling with LD555-TTZ (single labeling), LD655-TTZ (single labeling), or LD555-TTZ/LD655-TTZ (dual labeling), following the established viral labeling protocol. Excess unreacted dye was removed by sucrose density gradient centrifugation (5%, 10%, 15%, 20%, and 40% layers), and the viral fraction collected from above the 40% sucrose layer was used for spectral analysis.

Fluorescence excitation and emission spectra were recorded using a Horiba FluoroMax spectrofluorometer (FluoroMax Plus-C-SP). Excitation spectra were acquired over the visible wavelength ranges (450–650 nm for LD555-TTZ and 550–700 nm for LD655-TTZ) at fixed emission wavelengths of 569 nm and 669 nm, respectively. Emission spectra were recorded over the ranges (550-700 nm for LD555-TTZ and 650-800 nm for LD655-TTZ) for each sample upon excitation at 532 nm or 640 nm. The intensity was normalized to the maximum value in each spectrum for comparison. The resulting data were analyzed and plotted using GraphPad Prism software. No spectral shifts were observed between the free LD555 or LD655 dyes and their conjugated forms on HIV-1 viral particles.

### smFRET data acquisition

All smFRET experiments of Env on intact HIV-1 virions were performed using a customized prism-based total internal reflection fluorescence (prism-TIRF) microscope, as previously described.^27,28,47^ Fluorescently labeled HIV-1 virions were incubated in the absence or presence of 0.1 mg/mL ligands or antibodies in imaging buffer containing 50 mM Tris (pH 7.4), 50 mM NaCl, a triplet-state quencher cocktail, 2 mM protocatechuic acid (PCA), and 8 nM protocatechuate-3,4-dioxygenase (PCD) at room temperature for 30 minutes prior to imaging. Ligand and antibody concentrations were approximately 5-fold above their 95% inhibitory concentration (5x IC95). Fluorescently labeled HIV-1 virions were immobilized on a PEG-passivated, biotinylated quartz-coverslip imaging chamber coated with streptavidin. Based on the refractive index difference between quartz and the aqueous buffer, an evanescent field was generated by total internal reflection of a 532 nm single-wavelength laser (Ventus, Laser Quantum) directed onto a prism. The donor fluorophore labeled on Env was excited by this evanescent TIRF field, and the resulting fluorescence from both donor and acceptor fluorophores was collected through a water-immersion Nikon objective (60×, NA 1.27). Emission signals were then separated using a MultiCam LS image splitter (Cairn Research) equipped with a dichroic filter (Chroma) and directed through ET590/50 and ET690/50 emission filters (Chroma) corresponding to donor and acceptor channels, respectively. Fluorescence signals were recorded simultaneously using two synchronized sCMOS cameras (Hamamatsu ORCA-Flash4.0 V3) at a frame rate of 25 Hz for 80 seconds. Where indicated, virions were pre-incubated with the appropriate ligand or antibody for 30 minutes at room temperature before imaging.

### smFRET data processing and analysis

Data were viewed, processed, and analyzed by a customized SPARTAN software package^44^ and MATLAB-based scripts, as described previously by us and others.^17,23,24,27,28,47,59^ Image stacks from smFRET recordings (2,000 frames over 80 s) were extracted as individual fluorescence time trajectories (fluorescence traces) corresponding to donor and acceptor signals from labeled HIV-1 virions. At the single-molecule level, background fluorescence was estimated and subtracted based on the signal intensity at single-step photobleaching points. The FRET efficiency (FRET values or FRET in graphs) was calculated according to: FRET = I_A_/(I_D_+γI_A_), where I_D_ and I_A_ represent fluorescence intensities of the donor and acceptor, respectively, and the correlation coefficient γ compensates for crosstalk and differences in detection efficiency between two channels. The resulting FRET traces, time-resolved donor-to-acceptor energy transfer trajectories, reflect dynamic distance changes between the fluorophores, corresponding to real-time conformational dynamics of Env in the context of intact virions. To ensure data quality, stringent filtering criteria were applied. Fluorescence traces were automatically excluded if signals from either donor or acceptor were missing, if multiple fluorophores were present, or if signal-to-noise ratios were insufficient. Remaining traces were manually viewed to confirm anti-correlation between donor and acceptor intensities, a hallmark of genuine FRET events reflecting conformational transitions of active Env molecules. Only traces exhibiting this anti-correlation and consistent single-protomer labeling were retained for further analysis.

Accepted FRET traces were compiled into FRET histograms (conformational ensembles), representing the distribution of Env conformational states across multiple virions. Each histogram represents the mean ± SEM, determined from three randomly assigned subsets of FRET traces under identical experimental conditions. The number of FRET states was inferred by combining visual inspection of FRET trajectories with iterative Hidden Markov Modeling (HMM).^62,63^ Model initialization was guided by visually identified transitions and optimized through iterative segmentation and semi-automatic parameter refinement. Statistical evaluation of model fits indicated that a three-state model provided the simplest and most accurate representation of the data, yielding lower log-likelihood values than a two-state model and avoiding overfitting.

FRET histograms were further fitted to the sum of three Gaussian distributions using a least-squares fitting algorithm in MATLAB. Each Gaussian component represented a distinct conformational state of Env, and the area under each Gaussian estimated the relative occupancy of that state. Relative state occupancies are reported as mean ± SEM, derived from the fitted histograms. The corresponding fitting parameters are summarized in Table S1. The three FRET states correspond to distinct and reproducible conformational populations under different experimental conditions, consistent with previous Env smFRET studies employing different probe positions.^17,24,47,64^

Transition rates were derived from the above-mentioned FRET trace idealization, in which each FRET trajectory was converted into a time-correlated sequence of discrete states using a segmental *K*-means algorithm within an HMM framework.^62,63^ State-to-state transitions, indicating the locations and frequencies of conformational changes, were visualized as transition density plots (TDPs). Dwell time distributions, representing the duration a molecule remains in a specific conformational state before transitioning to another, were compiled into survival probability plots and fitted to a bi-exponential decay function (y = *A*_1_ exp*^−k1t^* + *A*_2_ exp*^−k2t^*). *A*_1_ and *A*_2_ are the amplitudes, and *k1* and *k2* are the corresponding rate constants. The overall transition rate was determined as the amplitude-weighted average of the two rate constants.

### Molecular dynamics simulation and analysis

Atomic coordinates of HIV-1 Env in the prefusion closed state were obtained by predicting the structure of the Env trimer using AlphaFold3. The input sequence consisted of three gp120 subunits (residues 30–505) and three gp41 subunits (residues 509–661) derived from HIV-1 BG505 Env (UniProt ID Q2N0S5). To enable site-specific fluorophore labeling, residues S398 in gp120 and R539 in gp41 of a single protomer were mutated to canonical lysine residues. The top-ranked predicted model showed acceptable agreement with the experimentally determined prefusion closed Env structure (RCSB PDB ID: 4TVP),^6^ with a backbone RMSD of 0.82 Å. Atomic coordinates of Env in the CD4-bound open state were based on the experimentally solved structure (RCSB PDB ID: 5VN3).^21^ Residues R542 (chain A) and T401 (chain J) were mutated to lysine residues using PyMOL (Schrödinger).

Atomic coordinates for the two fluorophores and linkers were generated with MarvinSketch (Chemaxon) and PyMOL. Force-field parameters for the fluorophores were generated using a fragment-based parameterization strategy. The fluorophores alone and model compounds consisting of the fluorophore covalently linked to a lysine side-chain fragment were both parameterized using GAFF/GAFF2, with AM1-BCC partial charges assigned using Antechamber. The latter was used to capture the local electronic environment introduced by covalent conjugation. During construction of the full fluorophore–protein conjugates, atom types and partial charges associated with the covalent linkage were assigned by transferring consistent parameter trends from the fluorophore–lysine model compounds, while preserving the original fluorophore parameters wherever applicable. The fluorophores were assembled with the protein, and the assembly was visualized in VMD. Final system preparation, including definition of covalent linkages and generation of topology and coordinate files, was performed using LEaP in AmberTools.

Each system was neutralized and solvated in explicit TIP3P water with periodic boundary conditions. Sodium and chloride ions were added to achieve a final salt concentration of 150 mM NaCl. Protein atoms were described using the Amber ff14SB force field.

Energy minimization was performed in multiple stages, followed by gradual heating from 0 to 300 K under constant volume (NVT) conditions. The systems were subsequently equilibrated under constant pressure (NPT) conditions at 1 atm to stabilize system density. Following equilibration, a pre-production simulation was carried out under NPT conditions for 200 ps with weak positional restraints applied to the protein backbone. Production simulations were then performed in the NPT ensemble at 300 K and 1 atm with all restraints removed. Atomic coordinates were saved every 5 ps. For each labeled system, three independent production trajectories of 10 ns were generated using different initial velocity seeds.

Dye positions sampled throughout the production trajectories were extracted for every simulation frame. All trajectory frames were structurally aligned to the protein backbone. The protein structure from a reference frame was retained, while fluorophore coordinates from all frames were superimposed to generate a composite dye ensemble representation.

Inter-dye distances were calculated on a per-frame basis using the center-of-mass (COM) coordinates of the fluorophore chromophores. Distance time traces were obtained directly from the production trajectories and smoothed using a centered moving average with a time window of 0.4 ns.

## Data availability

Data supporting the findings of this study are available within the paper and its supplementary information files.

## Acknowledgements

The authors thank Yuanyun Ao for the valuable discussions and contributions during the early stages of this study. The following reagent was obtained through BEI Resources, NIAID, NIH: Monoclonal Anti-Human Immunodeficiency Virus Type 1 (HIV-1) gp120 (PG16), ARP-12150. Research reported in this publication was supported by the National Institute of Allergy and Infectious Diseases of the National Institutes of Health (NIH) under Award Number R01AI181600 to M.L. The content is solely the responsibility of the authors and does not necessarily represent the official views of the NIH. This research was also supported in part by the Intramural Research Program of the NIH. The contributions of the NIH author were made as part of his official duties as a NIH federal employee, are in compliance with agency policy requirements, and are considered Works of the United States Government. However, the findings and conclusions presented in this paper are those of the author and do not necessarily reflect the views of the NIH or the U.S. Department of Health and Human Services.

## Author contributions

M.L. conceived the study. M.L. and W.X. designed the experiments. W.X. and N.K.G. performed mutagenesis, virus infectivity and neutralization assays, Western blotting, virus production, click-chemistry–based fluorescence labeling, and virus purification. M.L., Y.H., W.X., and R.K. conducted smFRET experiments and data analysis. J.L. performed molecular dynamics simulations. R.K., W.X., and N.K.G. performed fluorescence spectral measurements. H.B.A. and Y.H. assisted with cell maintenance and plasmid preparation. B. H. and R. W. conducted the Cryo-ET and negative staining analyses. B.Z., J.Y., D.D.H., P.A., and P.D.K. provided antibodies. M.L. and W.X. prepared the figures. M.L. and W.X. wrote the manuscript with input from all authors, especially from P.D.K. All authors reviewed and approved the final manuscript.

---

**Table S1.** Model-fitting statistics and parameters for FRET histograms. FRET histograms acquired from two structural perspectives (top: S401 – R542; bottom: N136 – S401) of Ab-incubated Env on native virions were fitted independently using constrained three-state models. Each state was described by a Gaussian distribution *N* (*µ*, σ^2^) corresponding to PT, PC, and CO states. The mean (*µ*) and standard deviation (*σ*) parameters were determined separately for each perspective based on visual inspection of trajectories exhibiting state-to-state transitions and idealization of individual trajectories using multi-state Hidden Markov modeling. State probabilities are reported as mean ± s.e.m. Goodness-of-fit was assessed using *R*^2^ and RMSE (root-mean-square error), with *R*^2^ values approaching 1 and RMSE values approaching 0 indicating a high-quality fit.

| Conformational populations of Ab-incubated Env on native virions | Curve fitting $R^2$ | RMSE | PT State | PC State | CO State |
| --- | --- | --- | --- | --- | --- |
| | | | $\mu$: 0.55 ± 0.05 | $\mu$: 0.28 ± 0.04 | $\mu$: 0.10 ± 0.02 |
| | | | $\sigma$: 0.15 ± 0.01 | $\sigma$: 0.09 ± 0.02 | $\sigma$: 0.07 ± 0.01 |
| Env<sub>BG505</sub> | 0.9732 | 8.427e-4 | 59% ± 15% | 24% ± 14% | 17% ± 6% |
| Env<sub>BG505</sub> + sCD4 + 17b | 0.9878 | 8.447e-4 | 28% ± 13% | 30% ± 13% | 42% ± 7% |
| Env<sub>BG505</sub> + 8ANC195 | 0.9734 | 9.605e-4 | 45% ± 14% | 32% ± 15% | 23% ± 9% |
| Env<sub>BG505</sub> + VRC34 | 0.9893 | 6.638e-4 | 34% ± 11% | 42% ± 15% | 24% ± 9% |
| Env<sub>BG505</sub> + PGT151 | 0.9895 | 6.858e-4 | 34% ± 14% | 47% ± 14% | 19% ± 5% |
| Env<sub>BG505</sub> + DH511.2_K3 | 0.9890 | 7.149e-4 | 36% ± 14% | 45% ± 14% | 19% ± 6% |
| Env<sub>BG505</sub> + VRC42 | 0.9865 | 8.237e-4 | 22% ± 10% | 53% ± 11% | 25% ± 5% |
| Env<sub>BG505</sub> + 10E8.4 | 0.9877 | 8.375e-4 | 25% ± 16% | 51% ± 16% | 24% ± 8% |
| Env<sub>BG505</sub> + iMab | 0.9756 | 8.503e-4 | 59% ± 16% | 27% ± 17% | 14% ± 11% |
| Env<sub>BG505</sub> + 10E8.4/iMab | 0.9700 | 0.0012 | 28% ± 10% | 35% ± 12% | 37% ± 14% |
| Env<sub>BG505</sub> + 10E8.4/iMab | 0.9700 | 0.0012 | 28% ± 10% | 35% ± 12% | 37% ± 14% |
| Env<sub>BG505</sub> + 10E8.4 + sCD4 + 17b | 0.9910 | 8.471e-4 | 23% ± 16% | 33% ± 14% | 44% ± 6% |

**Table S1. Model-fitting statistics and parameters for FRET histograms.**
| Conformational populations of Ab-incubated Env on native virions | Curve fitting $R^2$ | RMSE | PT State | PC State | CO State |
| --- | --- | --- | --- | --- | --- |
| | | | $\mu$: 0.10 ± 0.02 | $\mu$: 0.60 ± 0.01 | $\mu$: 0.32 ± 0.03 |
| | | | $\sigma$: 0.08 ± 0.01 | $\sigma$: 0.15 ± 0.01 | $\sigma$: 0.11 ± 0.01 |
| Env<sub>BG505</sub> | 0.9879 | 9.715e-4 | 42% ± 6% | 23% ± 11% | 35% ± 11% |
| Env<sub>BG505</sub> + sCD4 + 17b | 0.9841 | 8.701e-4 | 29% ± 8% | 26% ± 13% | 45% ± 15% |

**Table S2.**
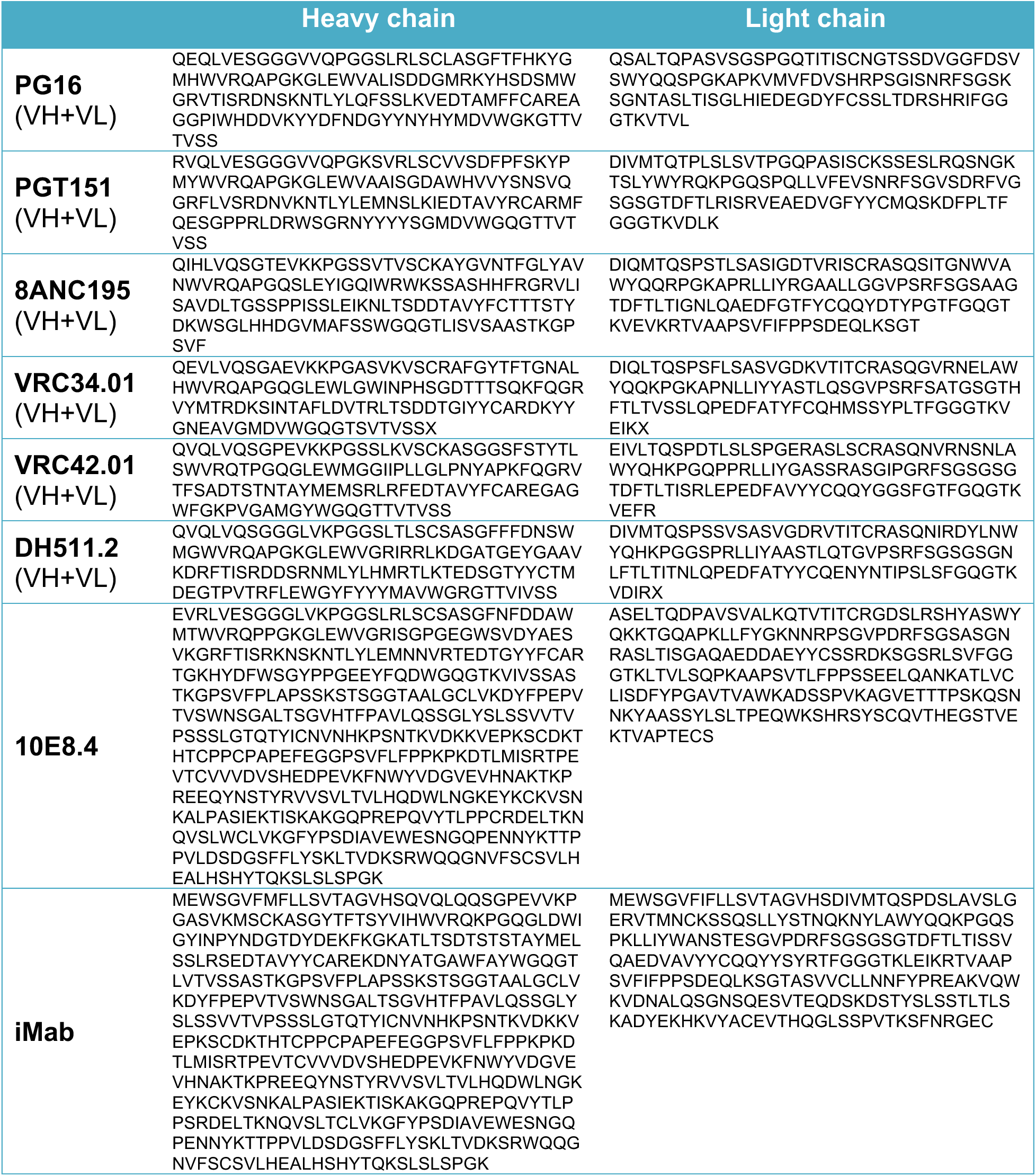
Amino acid sequences of heavy and light chains of antibodies used in this study.

**Fig. S1.**
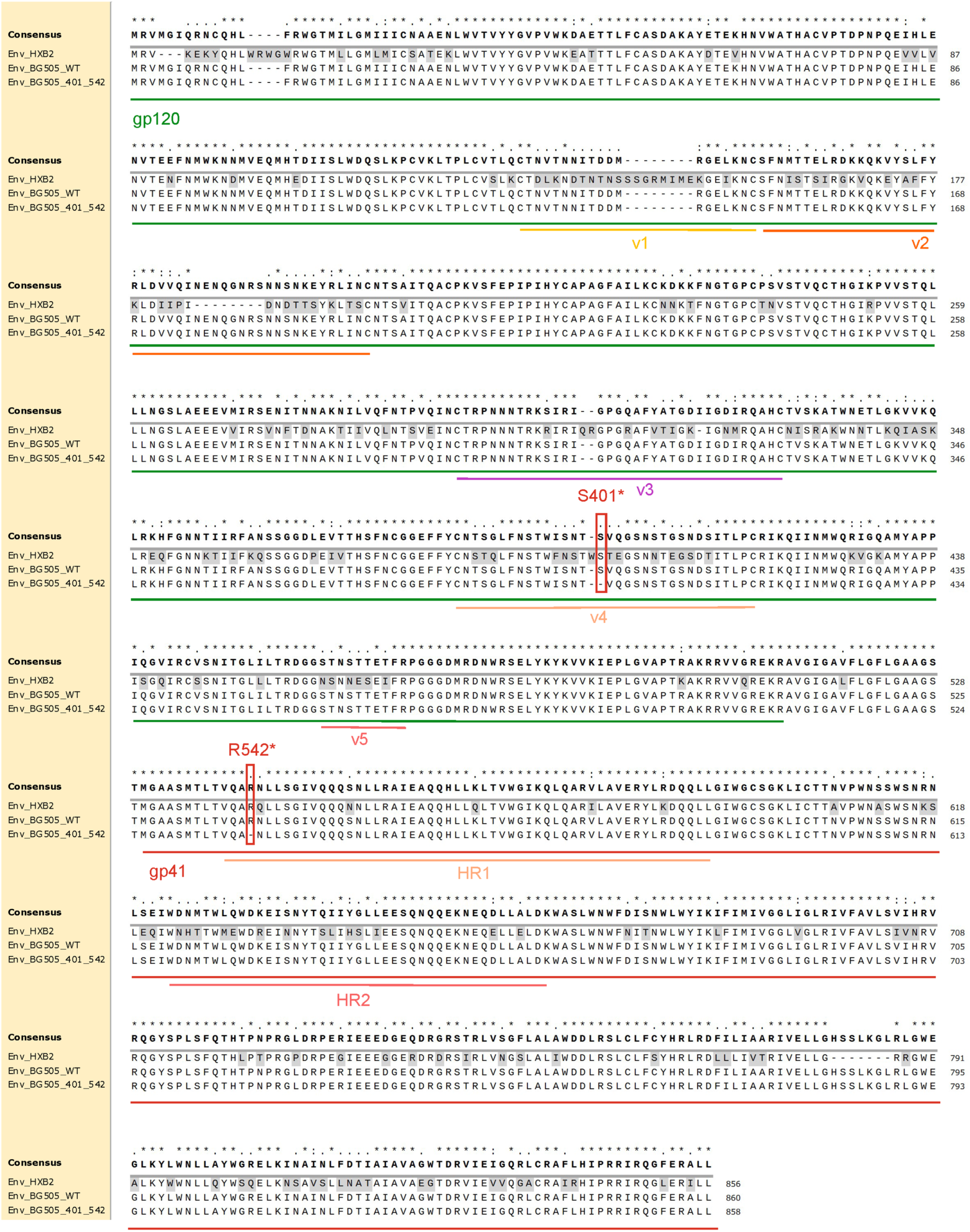
Primary structures of Env with a pair of dye-attached positions indicated on BG505 for smFRET studies.

**Figure S2.**
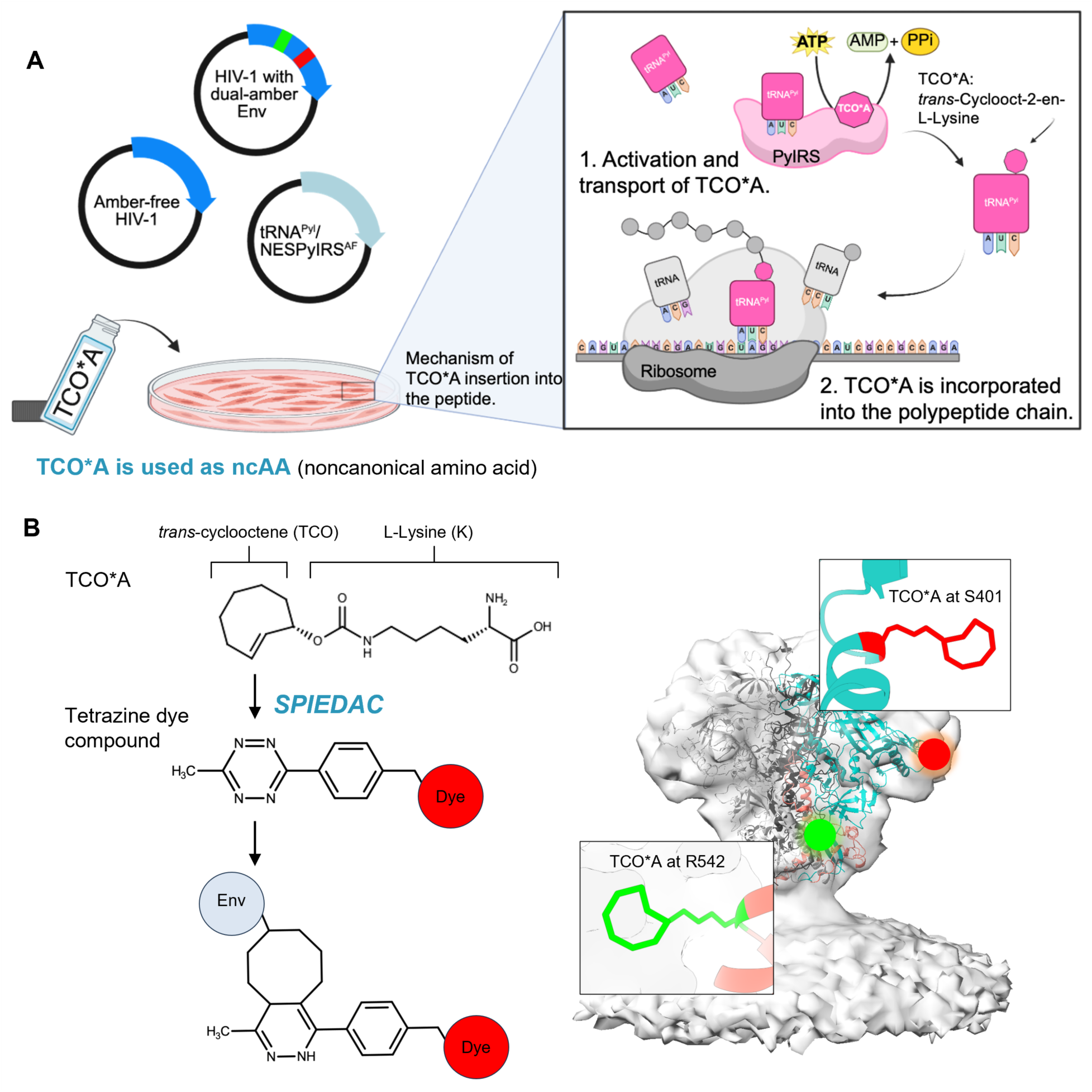
Amber suppression and click-labeling strategy for preparing HIV-1_Q23_ Env_BG505_ virions for smFRET. (**A**) Genetic code expansion via amber suppression. Amber suppressor tRNA and tRNA synthetase (tRNA^Pyl^/NESPylRS^AF^) incorporate the ncAA trans-cyclooct-2-en-L-lysine (TCO*A) at TAG codons engineered into Env (S401^TAG^ in gp120 and R542^TAG^ in gp41) on intact virions produced in mammalian cells. The supply of TCO*A to transfected cells enables its incorporation at the designated positions, providing reactive handles for subsequent fluorophore conjugation. (**B**) Bioorthogonal click labeling via SPIEDAC. Tetrazine-conjugated Cy3 and Cy5 derivatives (LD555-TTZ and LD655-TTZ) react with the strained alkene of TCO*A through strain-promoted inverse electron-demand Diels–Alder cycloaddition (SPIEDAC). The conjugated fluorophores (dyes) are depicted as red spheres. TCO*A Functional groups are shown in the modeled membrane-present Env trimers (right panel).

**Figure S3.**
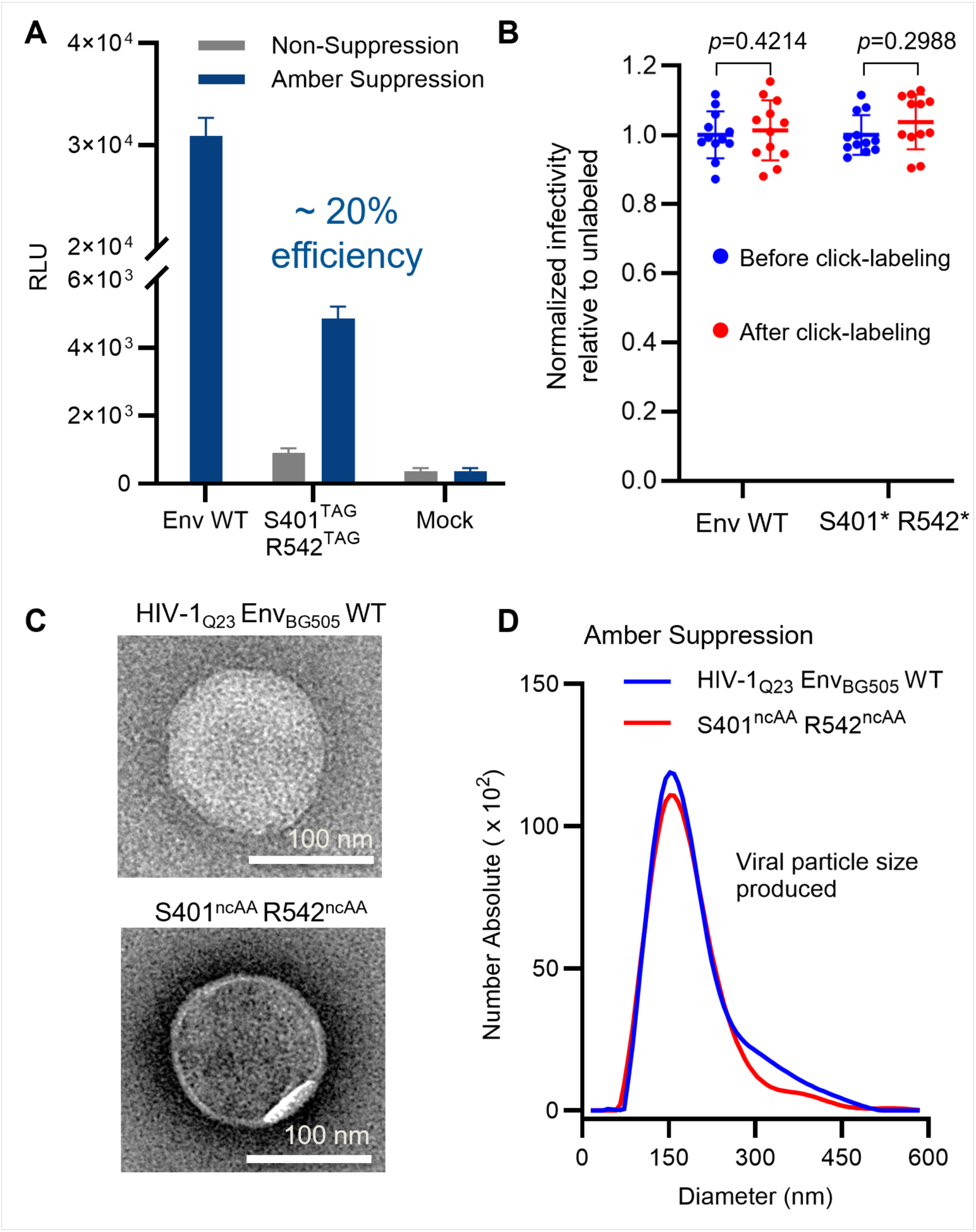
Dual-amber suppression efficiency of HIV-1_Q23_ Env_BG505_ S401^TAG^ R542^TAG^, infectivity before and after labeling, and particle size distribution of the resulting virions. (**A**) Infectivity of dual-ncAA (dual-amber) virions was measured on TZM-bl cells. Virions were produced by transfecting HEK293T cells with plasmids amber-free HIV-1_Q23_ Env_BG505_ S401^TAG^ R542^TAG^. Infectivity (mean ± SD, n=3) was normalized to wild-type Env_BG505_ (WT) under suppression conditions. With amber suppression (tRNA^Pyl^/NESPylRS^AF^ + TCO*A), the dual-amber construct retained ~20% of WT infectivity, whereas non-suppression controls (no supplement) showed negligible infectivity. Mock denotes particles produced without Env plasmids. (**B**) No significant difference in infectivity was detected before and after fluorescent labeling (S401* R542*) of clickable HIV-1_Q23_ Env_BG505_ S401^ncAA^ R542^ncAA^. *P* values were calculated using an unpaired two-tailed Welch’s t-test. (**C**) Example negative-staining transmission electron microscopy images of the resulting HIV-1_Q23_ Env_BG505_ S401^ncAA^ R542^ncAA^ virions produced under amber suppression conditions. (**D**) Particle size distributions of HIV-1 particles (as in panel B) measured by nanoparticle tracking analysis (NTA, ZetaView). Dual-ncAA Env virions exhibited size distributions comparable to WT Env virions.

**Figure S4.**
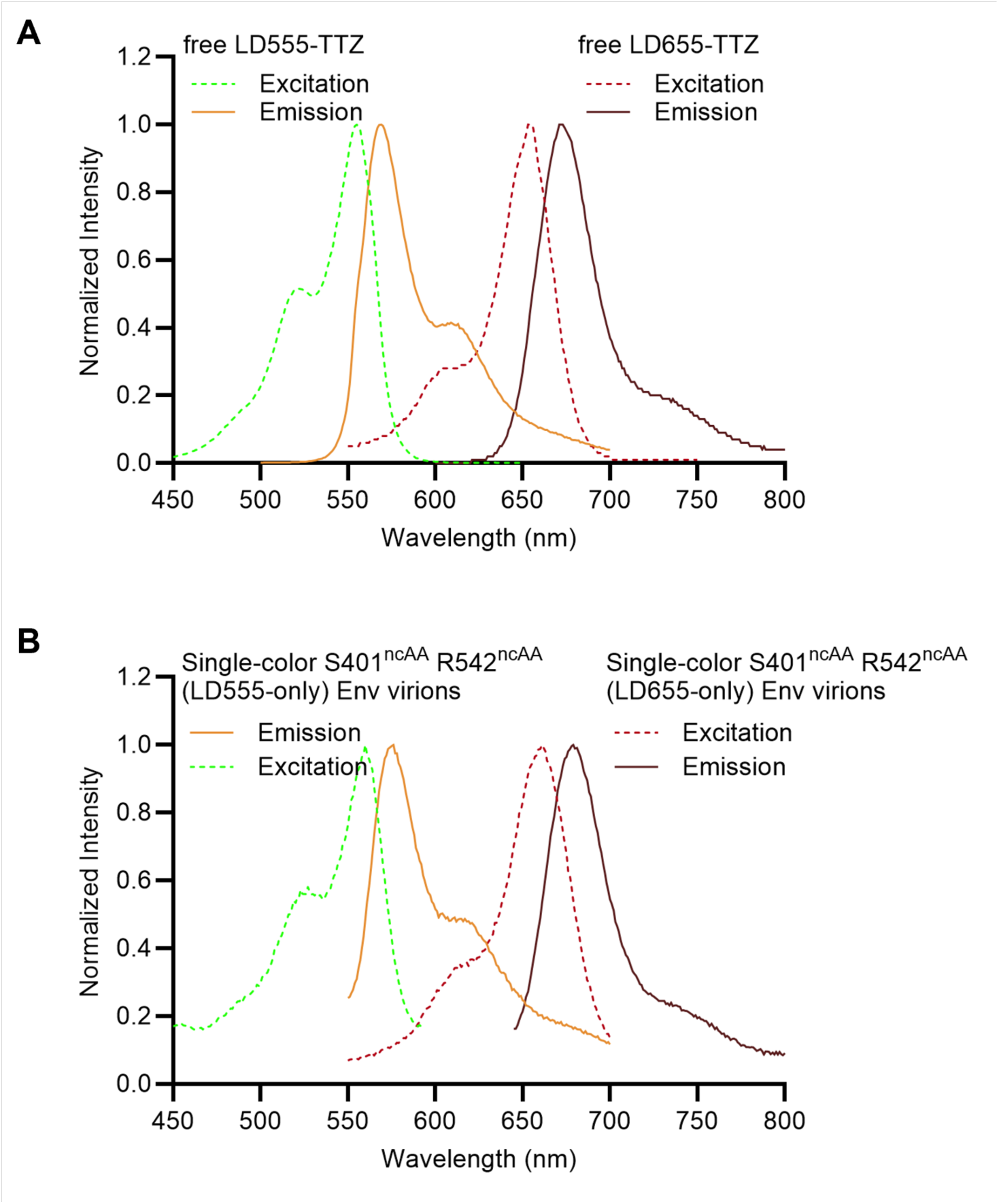
Biophysical fluorescence properties of functionalized fluorophores and single–color–labeled Env_BG505_ HIV-1_Q23_ S401^ncAA^ R542^ncAA^ virions. (**A**) Excitation and emission spectra of free LD555-tetrazine (LD555-TTZ) and LD655-tetrazine (LD655-TTZ). (**B**) Excitation and emission spectra of single-color–labeled S401^ncAA^ R542^ncAA^ virions, in which both ncAA sites (S401 on gp120 and R542 on gp41) were labeled with the same fluorophore via SPIEDAC click chemistry. LD555-only denotes virions labeled with LD555-TTZ at both sites, and LD655-only denotes virions labeled with LD655-TTZ at both sites. These single-color–labeled virions serve as spectral controls.

**Figure S5.**
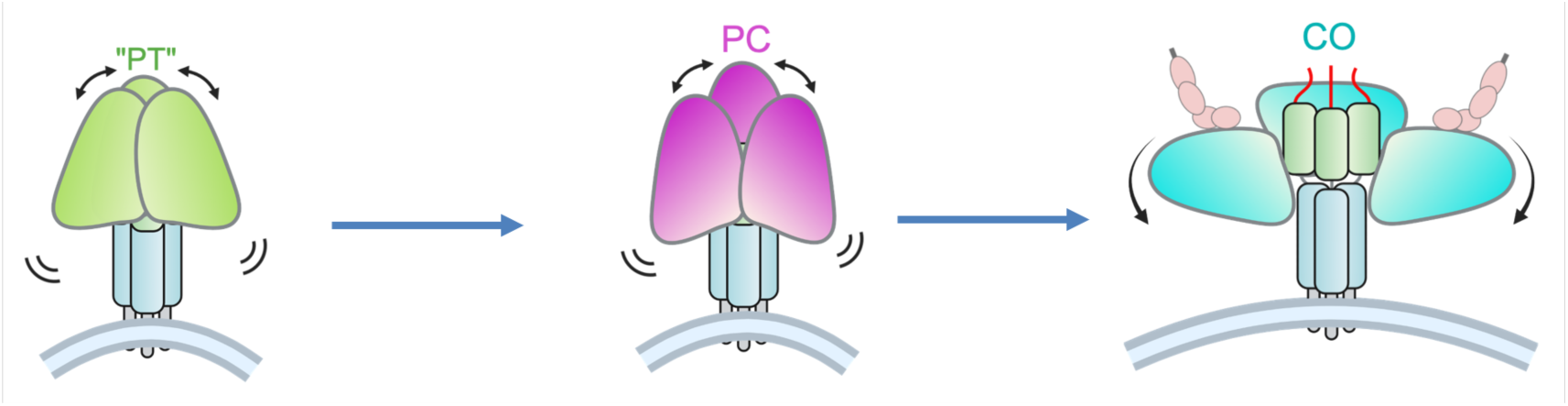
Relative protomer movements in symmetric PT, PC, and CO Env trimers. Cartoon representations of Env trimers in three major conformations: “PT” (pre-triggered, structurally unknown), PC (prefusion closed), and CO (CD4-bound open). This figure complements Figure 2A by showing the three conformations as symmetric trimers, whereas Figure 2A depicts a single protomer transitioning within an otherwise PT-state trimer.

**Figure S6.**
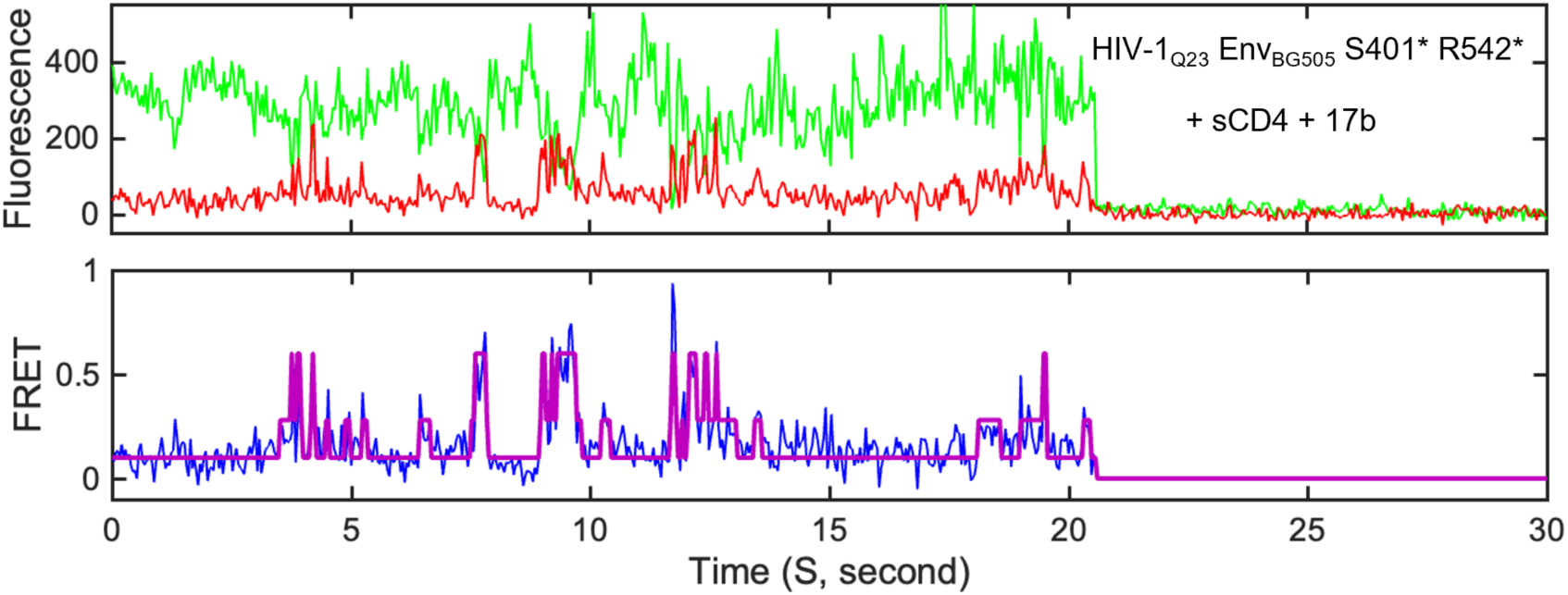
Representative traces of Env predominantly in the open conformation. Example fluorescence and FRET traces from an individual Env_BG505_ S401* R542* virion under opening conditions with soluble CD4 (sCD4) and the co-receptor–mimicking antibody 17b.

**Figure S7.**
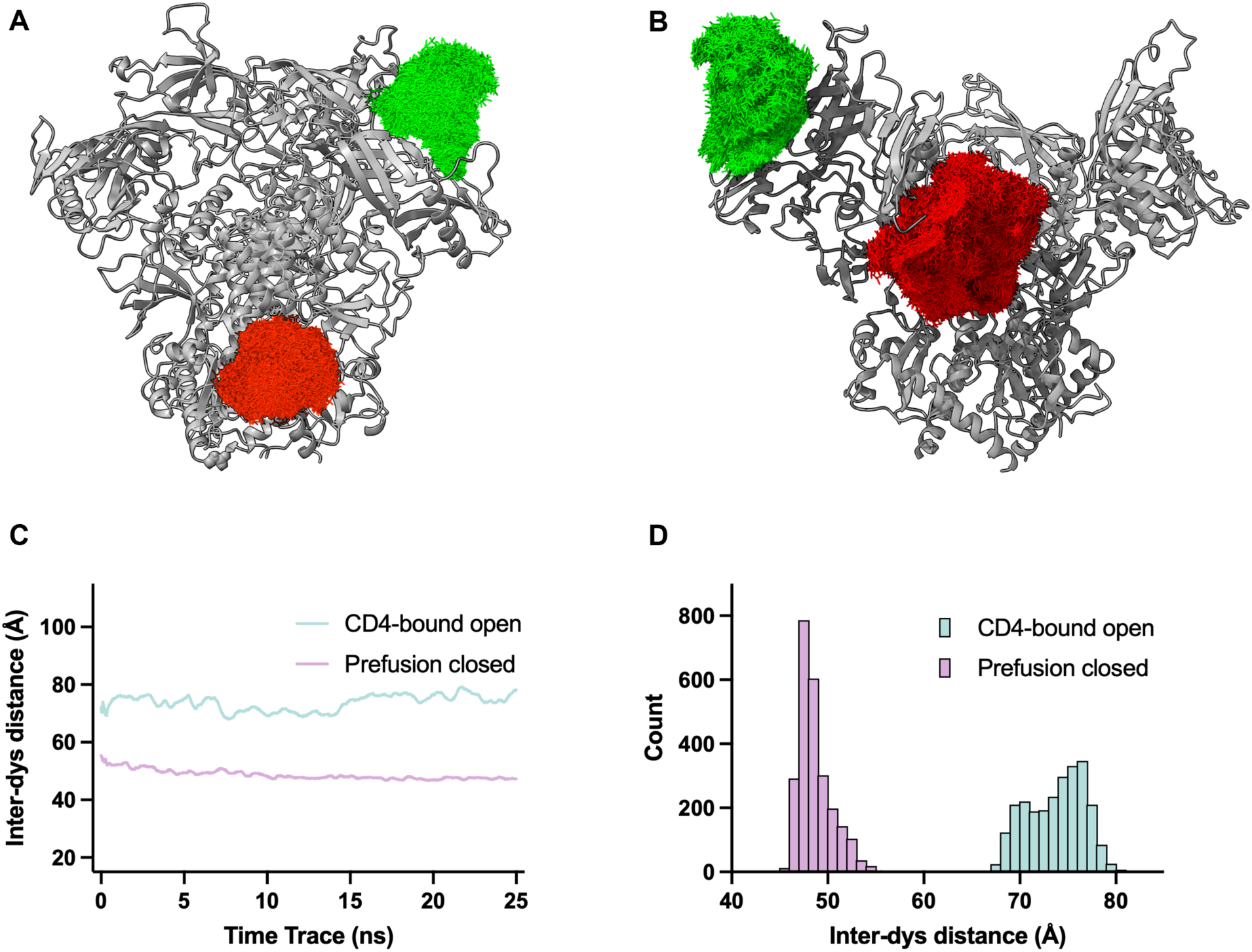
Molecular dynamics simulation of dye-conjugated soluble Env trimer in prefusion closed state and CD4-bound state. (**A**) Sampling space of dye conformations during a 10-ns molecular dynamics simulation of dye-conjugated soluble Env S401^TCO*-A^ – LD555 R542^TCO*A^ – LD655 in prefusion closed state (RCSB PDB ID: 4TVP). Dye positions were extracted from the equilibrated portion of the trajectory and superimposed after backbone alignment. LD555 is shown in green, LD655 in red, and the protein backbone in grey. (**B**) Sampling space of dye conformations during a 10-ns molecular dynamics simulation of dye-conjugated Env in CD4-bound open state (RCSB PDB ID: 5VN3). Dye positions were extracted and displayed as described in (A). (**C**) Inter-dye distance as a function of simulation time (25 ns) for dye-conjugated Env trimer in prefusion closed state and CD4-bound open state, calculated from the center-of-mass distance between the two dyes at each saved frame. Distance traces were smoothed using a centred moving average window of 40 ps for visualization. (**D**) Distribution of inter-dye distances. Distances were calculated as the center-of-mass separation between the two dyes over the trajectories shown in (C). Mean inter-dye distances are 48.68 ± 1.75 Å (prefusion closed state) and 73.75 ± 2.94 Å (CD4-bound open state), respectively (mean ± SD).

**Figure S8.**
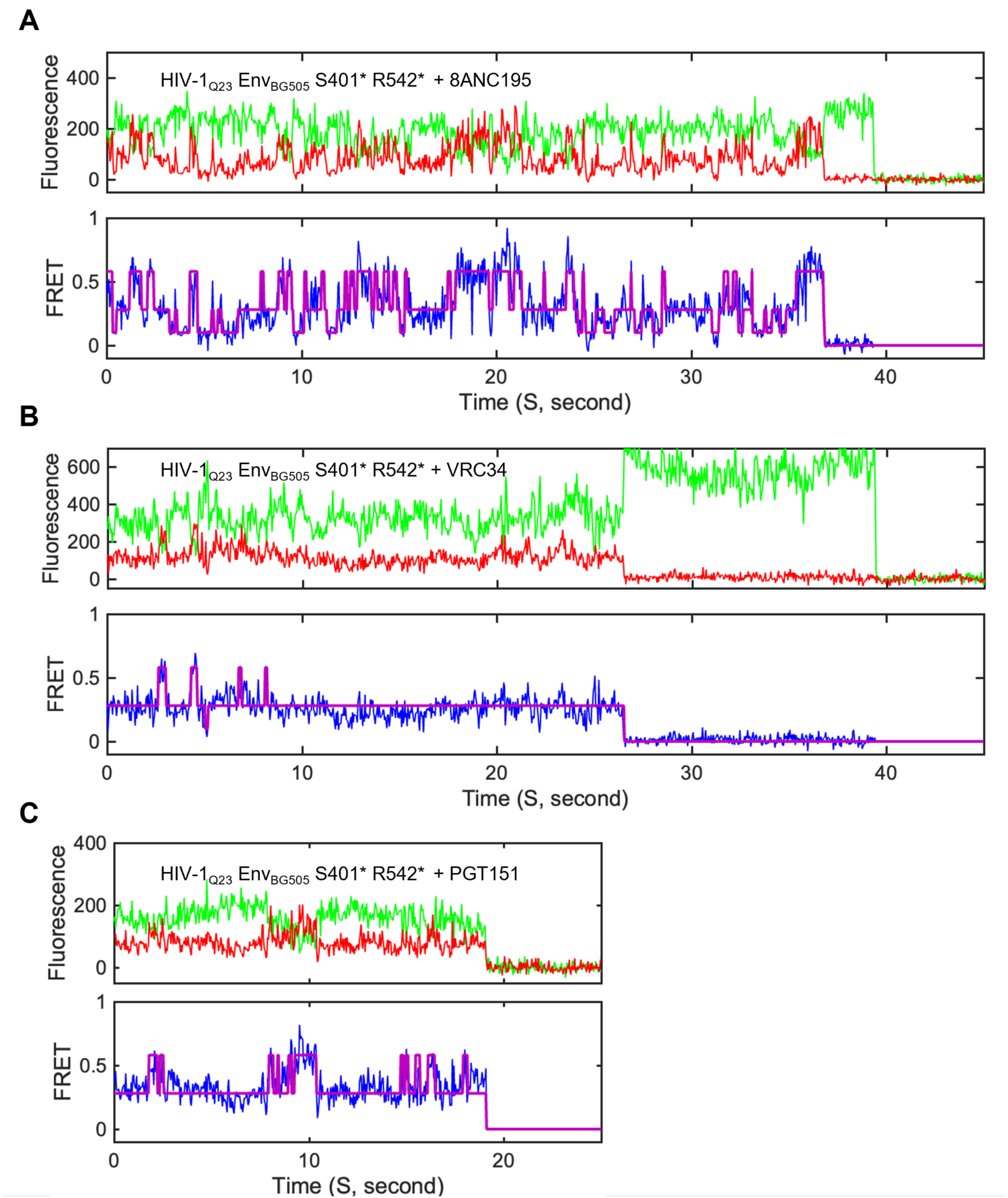
Representative smFRET traces of HIV-1Q23 EnvBG505 S401* R542* with gp120-gp41 interface or fusion peptide bNAbs. (**A – C**) Example donor and acceptor fluorescence (top) and corresponding FRET efficiency traces (bottom) from an individual EnvBG505 S401* R542* in the presence of 8ANC195 (**A**), VRC34 (**B**), and PGT151 (**C**).

**Figure S9.**
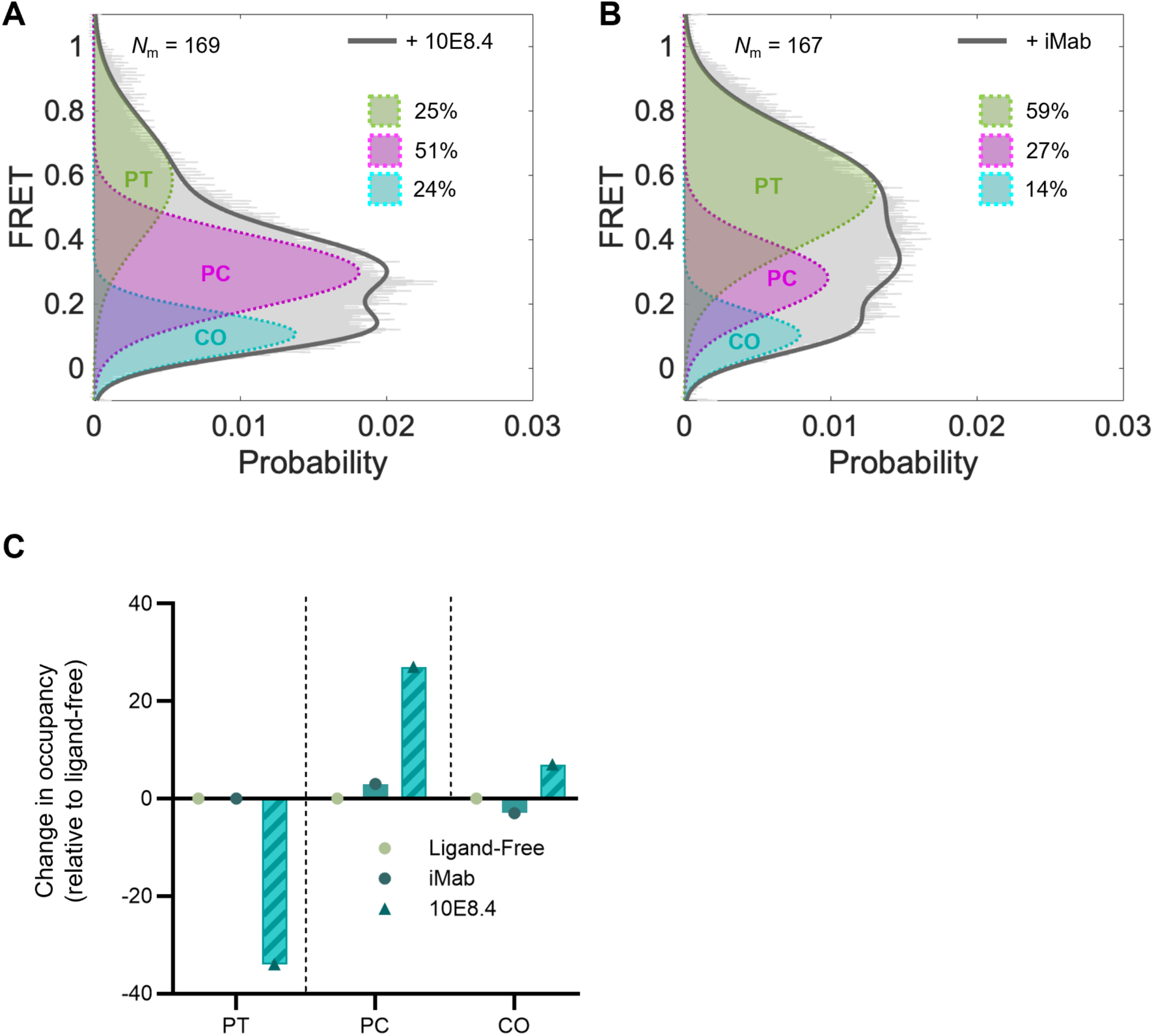
10E8.4 enriches Env in the PC state, while iMab itself has no effect on Env conformations. (**A, B**) FRET histograms of Env_BG505_ S401* R542* in the presence of 10E8.4 (**A**), or iMab (**B**). (**C**) Changes in state occupancy of Env induced by 10E8.4 or iMab, relative to the ligand-free condition.

**Figure S10.**
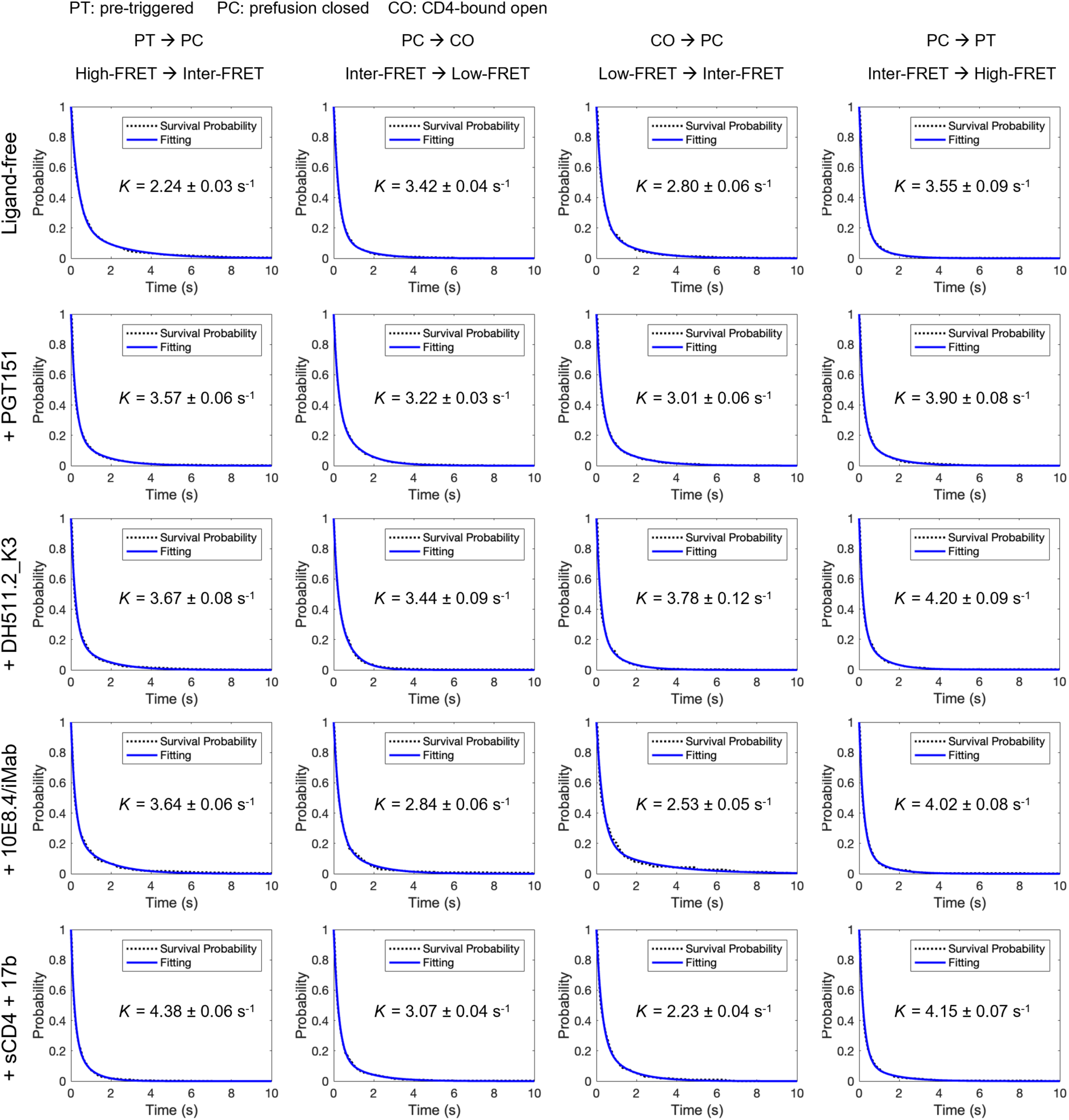
Survival probability plots used to estimate transition rates among Env conformations. Survival probability plots showing the probability of occupying a given FRET-defined conformation as a function of dwell time. Data points represent pooled dwell times from individual molecules, and curves show exponential fits used to extract transition rate constants (see Methods). These rate estimates correspond to the transitions summarized in Figure 5.

## References

1. Wyatt, R., and Sodroski, J. (1998). The HIV-1 envelope glycoproteins: fusogens, antigens, and immunogens. Science 280, 1884–1888. 10.1126/science.280.5371.1884.

2. Chen, B. (2019). Molecular Mechanism of HIV-1 Entry. Trends Microbiol 27, 878–891. 10.1016/j.tim.2019.06.002.

3. Wang, Q., Finzi, A., and Sodroski, J. (2020). The Conformational States of the HIV-1 Envelope Glycoproteins. Trends Microbiol 28, 655–667. 10.1016/j.tim.2020.03.007.

4. Klasse, P.J., Sanders, R.W., Ward, A.B., Wilson, I.A., and Moore, J.P. (2025). The HIV-1 envelope glycoprotein: structure, function and interactions with neutralizing antibodies. Nat Rev Microbiol 23, 734–752. 10.1038/s41579-025-01206-6.

5. Kwong, P.D., Wyatt, R., Robinson, J., Sweet, R.W., Sodroski, J., and Hendrickson, W.A. (1998). Structure of an HIV gp120 envelope glycoprotein in complex with the CD4 receptor and a neutralizing human antibody. Nature 393, 648–659. 10.1038/31405.

6. Pancera, M., Zhou, T., Druz, A., Georgiev, I.S., Soto, C., Gorman, J., Huang, J., Acharya, P., Chuang, G.Y., Ofek, G., et al. (2014). Structure and immune recognition of trimeric pre-fusion HIV-1 Env. Nature 514, 455–461. 10.1038/nature13808.

7. Lyumkis, D., Julien, J.P., de Val, N., Cupo, A., Potter, C.S., Klasse, P.J., Burton, D.R., Sanders, R.W., Moore, J.P., Carragher, B., et al. (2013). Cryo-EM structure of a fully glycosylated soluble cleaved HIV-1 envelope trimer. Science 342, 1484–1490. 10.1126/science.1245627.

8. Shaik, M.M., Peng, H., Lu, J., Rits-Volloch, S., Xu, C., Liao, M., and Chen, B. (2019). Structural basis of coreceptor recognition by HIV-1 envelope spike. Nature 565, 318–323. 10.1038/s41586-018-0804-9.

9. Yang, X., Kurteva, S., Ren, X., Lee, S., and Sodroski, J. (2005). Stoichiometry of envelope glycoprotein trimers in the entry of human immunodeficiency virus type 1. J Virol 79, 12132–12147. 10.1128/JVI.79.19.12132-12147.2005.

10. Yang, X., Kurteva, S., Ren, X., Lee, S., and Sodroski, J. (2006). Subunit stoichiometry of human immunodeficiency virus type 1 envelope glycoprotein trimers during virus entry into host cells. J Virol 80, 4388–4395. 10.1128/JVI.80.9.4388-4395.2006.

11. Sougrat, R., Bartesaghi, A., Lifson, J.D., Bennett, A.E., Bess, J.W., Zabransky, D.J., and Subramaniam, S. (2007). Electron tomography of the contact between T cells and SIV/HIV-1: Implications for viral entry. Plos Pathogens 3, 571–581. ARTN e63. 10.1371/journal.ppat.0030063.

12. Liu, J., Bartesaghi, A., Borgnia, M.J., Sapiro, G., and Subramaniam, S. (2008). Molecular architecture of native HIV-1 gp120 trimers. Nature 455, 109–113. 10.1038/nature07159.

13. Klasse, P.J. (2007). Modeling how many envelope glycoprotein trimers per virion participate in human immunodeficiency virus infectivity and its neutralization by antibody. Virology 369, 245–262. 10.1016/j.virol.2007.06.044.

14. Brandenberg, O.F., Magnus, C., Regoes, R.R., and Trkola, A. (2015). The HIV-1 Entry Process: A Stoichiometric View. Trends Microbiol 23, 763–774. 10.1016/j.tim.2015.09.003.

15. Brandenberg, O.F., Magnus, C., Rusert, P., Regoes, R.R., and Trkola, A. (2015). Different Infectivity of HIV-1 Strains Is Linked to Number of Envelope Trimers Required for Entry. Plos Pathogens 11. ARTN e1004595. 10.1371/journal.ppat.1004595.

16. Katte, R.H., Xu, W., Han, Y., Hong, X., and Lu, M. (2025). Inter-protomer opening cooperativity of envelope trimers positively correlates with HIV-1 entry stoichiometry. mBio 16, e0275424. 10.1128/mbio.02754-24.

17. Munro, J.B., Gorman, J., Ma, X., Zhou, Z., Arthos, J., Burton, D.R., Koff, W.C., Courter, J.R., Smith, A.B., 3rd, Kwong, P.D., et al. (2014). Conformational dynamics of single HIV-1 envelope trimers on the surface of native virions. Science 346, 759–763. 10.1126/science.1254426.

18. Munro, J.B., and Mothes, W. (2015). Structure and Dynamics of the Native HIV-1 Env Trimer. J Virol 89, 5752–5755. 10.1128/JVI.03187-14.

19. Herschhorn, A., Ma, X., Gu, C., Ventura, J.D., Castillo-Menendez, L., Melillo, B., Terry, D.S., Smith, A.B., 3rd, Blanchard, S.C., Munro, J.B., et al. (2016). Release of gp120 Restraints Leads to an Entry-Competent Intermediate State of the HIV-1 Envelope Glycoproteins. mBio 7, e01598–01516. 10.1128/mBio.01598-16.

20. Herschhorn, A., Gu, C., Moraca, F., Ma, X., Farrell, M., Smith, A.B., 3rd, Pancera, M., Kwong, P.D., Schon, A., Freire, E., et al. (2017). The beta20-beta21 of gp120 is a regulatory switch for HIV-1 Env conformational transitions. Nat Commun 8, 1049. 10.1038/s41467-017-01119-w.

21. Ozorowski, G., Pallesen, J., de Val, N., Lyumkis, D., Cottrell, C.A., Torres, J.L., Copps, J., Stanfield, R.L., Cupo, A., Pugach, P., et al. (2017). Open and closed structures reveal allostery and pliability in the HIV-1 envelope spike. Nature 547, 360–363. 10.1038/nature23010.

22. Henderson, R., Lu, M., Zhou, Y., Mu, Z., Parks, R., Han, Q., Hsu, A.L., Carter, E., Blanchard, S.C., Edwards, R.J., et al. (2020). Disruption of the HIV-1 Envelope allosteric network blocks CD4-induced rearrangements. Nat Commun 11, 520. 10.1038/s41467-019-14196-w.

23. Alsahafi, N., Bakouche, N., Kazemi, M., Richard, J., Ding, S., Bhattacharyya, S., Das, D., Anand, S.P., Prevost, J., Tolbert, W.D., et al. (2019). An Asymmetric Opening of HIV-1 Envelope Mediates Antibody-Dependent Cellular Cytotoxicity. Cell Host Microbe 25, 578–587 e575. 10.1016/j.chom.2019.03.002.

24. Ma, X., Lu, M., Gorman, J., Terry, D.S., Hong, X., Zhou, Z., Zhao, H., Altman, R.B., Arthos, J., Blanchard, S.C., et al. (2018). HIV-1 Env trimer opens through an asymmetric intermediate in which individual protomers adopt distinct conformations. Elife 7, e34271. 10.7554/eLife.34271.

25. Dam, K.A., Fan, C., Yang, Z., and Bjorkman, P.J. (2023). Intermediate conformations of CD4-bound HIV-1 Env heterotrimers. Nature 623, 1017–1025. 10.1038/s41586-023-06639-8.

26. Li, W., Qin, Z., Nand, E., Grunst, M.W., Grover, J.R., Bess, J.W., Jr., Lifson, J.D., Zwick, M.B., Tagare, H.D., Uchil, P.D., and Mothes, W. (2023). HIV-1 Env trimers asymmetrically engage CD4 receptors in membranes. Nature 623, 1026–1033. 10.1038/s41586-023-06762-6.

27. Lee, M., Lu, M., Zhang, B., Zhou, T., Katte, R., Han, Y., Rawi, R., and Kwong, P.D. (2024). HIV-1-envelope trimer transitions from prefusion-closed to CD4-bound-open conformations through an occluded-intermediate state. Comput Struct Biotechnol J 23, 4192–4204. 10.1016/j.csbj.2024.11.020.

28. Thakur, B., Katte, R.H., Xu, W., Janowska, K., Sammour, S., Henderson, R., Lu, M., Kwong, P.D., and Acharya, P. (2025). Conformational trajectory of the HIV-1 fusion peptide during CD4-induced envelope opening. Nat Commun 16, 4595. 10.1038/s41467-025-59721-2.

29. Ladinsky, M.S., Gnanapragasam, P.N., Yang, Z., West, A.P., Kay, M.S., and Bjorkman, P.J. (2020). Electron tomography visualization of HIV-1 fusion with target cells using fusion inhibitors to trap the pre-hairpin intermediate. Elife 9. 10.7554/eLife.58411.

30. Buzon, V., Natrajan, G., Schibli, D., Campelo, F., Kozlov, M.M., and Weissenhorn, W. (2010). Crystal structure of HIV-1 gp41 including both fusion peptide and membrane proximal external regions. PLoS Pathog 6, e1000880. 10.1371/journal.ppat.1000880.

31. Zhao, C., Li, H., Swartz, T.H., and Chen, B.K. (2022). The HIV Env Glycoprotein Conformational States on Cells and Viruses. mBio 13, e0182521. 10.1128/mbio.01825-21.

32. Kwong, P.D., Doyle, M.L., Casper, D.J., Cicala, C., Leavitt, S.A., Majeed, S., Steenbeke, T.D., Venturi, M., Chaiken, I., Fung, M., et al. (2002). HIV-1 evades antibody-mediated neutralization through conformational masking of receptor-binding sites. Nature 420, 678–682. 10.1038/nature01188.

33. Haynes, B.F., Burton, D.R., and Mascola, J.R. (2019). Multiple roles for HIV broadly neutralizing antibodies. Sci Transl Med 11. 10.1126/scitranslmed.aaz2686.

34. Sok, D., and Burton, D.R. (2018). Recent progress in broadly neutralizing antibodies to HIV. Nat Immunol 19, 1179–1188. 10.1038/s41590-018-0235-7.

35. Haynes, B.F., Wiehe, K., Borrow, P., Saunders, K.O., Korber, B., Wagh, K., McMichael, A.J., Kelsoe, G., Hahn, B.H., Alt, F., and Shaw, G.M. (2023). Strategies for HIV-1 vaccines that induce broadly neutralizing antibodies. Nat Rev Immunol 23, 142–158. 10.1038/s41577-022-00753-w.

36. Kwong, P.D., and Mascola, J.R. (2018). HIV-1 Vaccines Based on Antibody Identification, B Cell Ontogeny, and Epitope Structure. Immunity 48, 855–871. 10.1016/j.immuni.2018.04.029.

37. Ivan, B., Sun, Z., Subbaraman, H., Friedrich, N., and Trkola, A. (2019). CD4 occupancy triggers sequential pre-fusion conformational states of the HIV-1 envelope trimer with relevance for broadly neutralizing antibody activity. PLoS Biol 17, e3000114. 10.1371/journal.pbio.3000114.

38. Huang, Y., Yu, J., Lanzi, A., Yao, X., Andrews, C.D., Tsai, L., Gajjar, M.R., Sun, M., Seaman, M.S., Padte, N.N., and Ho, D.D. (2016). Engineered Bispecific Antibodies with Exquisite HIV-1-Neutralizing Activity. Cell 165, 1621–1631. 10.1016/j.cell.2016.05.024.

39. Padte, N.N., Yu, J., Huang, Y., and Ho, D.D. (2018). Engineering multi-specific antibodies against HIV-1. Retrovirology 15, 60. 10.1186/s12977-018-0439-9.

40. Nikic, I., Kang, J.H., Girona, G.E., Aramburu, I.V., and Lemke, E.A. (2015). Labeling proteins on live mammalian cells using click chemistry. Nat Protoc 10, 780–791. 10.1038/nprot.2015.045.

41. Sakin, V., Hanne, J., Dunder, J., Anders-Osswein, M., Laketa, V., Nikic, I., Krausslich, H.G., Lemke, E.A., and Muller, B. (2017). A Versatile Tool for Live-Cell Imaging and Super-Resolution Nanoscopy Studies of HIV-1 Env Distribution and Mobility. Cell Chem Biol 24, 635–645 e635. 10.1016/j.chembiol.2017.04.007.

42. Xu, W., Gonepudi, N.K., and Lu, M. (2025). Protocol for click labeling of HIV-1 envelope on amber-free virions prepared using genetic code expansion. STAR Protoc 6, 103995. 10.1016/j.xpro.2025.103995.

43. Roy, R., Hohng, S., and Ha, T. (2008). A practical guide to single-molecule FRET. Nat Methods 5, 507–516. 10.1038/Nmeth.1208.

44. Juette, M.F., Terry, D.S., Wasserman, M.R., Altman, R.B., Zhou, Z., Zhao, H., and Blanchard, S.C. (2016). Single-molecule imaging of non-equilibrium molecular ensembles on the millisecond timescale. Nat Methods 13, 341–344. 10.1038/nmeth.3769.

45. Lerner, E., Cordes, T., Ingargiola, A., Alhadid, Y., Chung, S., Michalet, X., and Weiss, S. (2018). Toward dynamic structural biology: Two decades of single-molecule Forster resonance energy transfer. Science 359, eaan1133. 10.1126/science.aan1133.

46. Lu, M., Ma, X., and Mothes, W. (2019). Illuminating the virus life cycle with single-molecule FRET imaging. Adv Virus Res 105, 239–273. 10.1016/bs.aivir.2019.07.004.

47. Ao, Y., Grover, J.R., Gifford, L., Han, Y., Zhong, G., Katte, R., Li, W., Bhattacharjee, R., Zhang, B., Sauve, S., et al. (2024). Bioorthogonal click labeling of an amber-free HIV-1 provirus for in-virus single molecule imaging. Cell Chem Biol 31, 487–501 e487. 10.1016/j.chembiol.2023.12.017.

48. Kwon, Y.D., Pancera, M., Acharya, P., Georgiev, I.S., Crooks, E.T., Gorman, J., Joyce, M.G., Guttman, M., Ma, X., Narpala, S., et al. (2015). Crystal structure, conformational fixation and entry-related interactions of mature ligand-free HIV-1 Env. Nat Struct Mol Biol 22, 522–531. 10.1038/nsmb.3051.

49. Li, Z., Li, W., Lu, M., Bess, J., Jr., Chao, C.W., Gorman, J., Terry, D.S., Zhang, B., Zhou, T., Blanchard, S.C., et al. (2020). Subnanometer structures of HIV-1 envelope trimers on aldrithiol-2-inactivated virus particles. Nat Struct Mol Biol 27, 726–734. 10.1038/s41594-020-0452-2.

50. Nikic, I., Estrada Girona, G., Kang, J.H., Paci, G., Mikhaleva, S., Koehler, C., Shymanska, N.V., Ventura Santos, C., Spitz, D., and Lemke, E.A. (2016). Debugging Eukaryotic Genetic Code Expansion for Site-Specific Click-PAINT Super-Resolution Microscopy. Angew Chem Int Ed Engl 55, 16172–16176. 10.1002/anie.201608284.

51. Plass, T., Milles, S., Koehler, C., Schultz, C., and Lemke, E.A. (2011). Genetically encoded copper-free click chemistry. Angew Chem Int Ed Engl 50, 3878–3881. 10.1002/anie.201008178.

52. Plass, T., Milles, S., Koehler, C., Szymanski, J., Mueller, R., Wiessler, M., Schultz, C., and Lemke, E.A. (2012). Amino acids for Diels-Alder reactions in living cells. Angew Chem Int Ed Engl 51, 4166–4170. 10.1002/anie.201108231.

53. Blackman, M.L., Royzen, M., and Fox, J.M. (2008). Tetrazine ligation: fast bioconjugation based on inverse-electron-demand Diels-Alder reactivity. J Am Chem Soc 130, 13518–13519. 10.1021/ja8053805.

54. Nikic, I., and Lemke, E.A. (2015). Genetic code expansion enabled site-specific dual-color protein labeling: superresolution microscopy and beyond. Curr Opin Chem Biol 28, 164–173. 10.1016/j.cbpa.2015.07.021.

55. Gonepudi, N.K., Xu, W., Awuah, H.B., and Lu, M. (2026). Bioorthogonal click-chemistry labeling and visualization of dual noncanonical amino acid-incorporated HIV-1 Env on intact virions. Methods Enzymol 731, 167–191. 10.1016/bs.mie.2026.05.005.

56. Xu, W., Gonepudi, N.K., Han, Y., and Lu, M. (2026). Genetic code expansion for single noncanonical amino acid incorporation into SARS-CoV-2 Omicron spike on amber-free lentiviral particles. Methods Enzymol 731, 141–165. 10.1016/bs.mie.2026.05.022.

57. Walker, L.M., Phogat, S.K., Chan-Hui, P.Y., Wagner, D., Phung, P., Goss, J.L., Wrin, T., Simek, M.D., Fling, S., Mitcham, J.L., et al. (2009). Broad and potent neutralizing antibodies from an African donor reveal a new HIV-1 vaccine target. Science 326, 285–289. 10.1126/science.1178746.

58. Blattner, C., Lee, J.H., Sliepen, K., Derking, R., Falkowska, E., de la Pena, A.T., Cupo, A., Julien, J.P., van Gils, M., Lee, P.S., et al. (2014). Structural delineation of a quaternary, cleavage-dependent epitope at the gp41-gp120 interface on intact HIV-1 Env trimers. Immunity 40, 669–680. 10.1016/j.immuni.2014.04.008.

59. Das, D.K., Govindan, R., Nikic-Spiegel, I., Krammer, F., Lemke, E.A., and Munro, J.B. (2018). Direct Visualization of the Conformational Dynamics of Single Influenza Hemagglutinin Trimers. Cell 174, 926–937 e912. 10.1016/j.cell.2018.05.050.

60. Das, D.K., Bulow, U., Diehl, W.E., Durham, N.D., Senjobe, F., Chandran, K., Luban, J., and Munro, J.B. (2020). Conformational changes in the Ebola virus membrane fusion machine induced by pH, Ca2^+^, and receptor binding. PLoS Biol 18, e3000626. 10.1371/journal.pbio.3000626.

61. Lu, M., Uchil, P.D., Li, W., Zheng, D., Terry, D.S., Gorman, J., Shi, W., Zhang, B., Zhou, T., Ding, S., et al. (2020). Real-Time Conformational Dynamics of SARS-CoV-2 Spikes on Virus Particles. Cell Host Microbe 28, 880–891 e888. 10.1016/j.chom.2020.11.001.

62. McKinney, S.A., Joo, C., and Ha, T. (2006). Analysis of single-molecule FRET trajectories using hidden Markov modeling. Biophys J 91, 1941–1951. 10.1529/biophysj.106.082487.

63. Qin, F. (2004). Restoration of single-channel currents using the segmental k-means method based on hidden Markov modeling. Biophys J 86, 1488–1501. 10.1016/S0006-3495(04)74217-4.

64. Lu, M. (2021). Single-Molecule FRET Imaging of Virus Spike-Host Interactions. Viruses 13, 332. 10.3390/v13020332.

65. Leonhardt, S.A., Purdy, M.D., Grover, J.R., Yang, Z., Poulos, S., McIntire, W.E., Tatham, E.A., Erramilli, S.K., Nosol, K., Lai, K.K., et al. (2023). Antiviral HIV-1 SERINC restriction factors disrupt virus membrane asymmetry. Nat Commun 14, 4368. 10.1038/s41467-023-39262-2.

66. Lu, M., Ma, X., Castillo-Menendez, L.R., Gorman, J., Alsahafi, N., Ermel, U., Terry, D.S., Chambers, M., Peng, D., Zhang, B., et al. (2019). Associating HIV-1 envelope glycoprotein structures with states on the virus observed by smFRET. Nature 568, 415–419. 10.1038/s41586-019-1101-y.

67. Richard, J., Grunst, M.W., Niu, L., Diaz-Salinas, M.A., Zhu, L., Tolbert, W.D., Marchitto, L., Zhou, F., Bourassa, C., Kim, H., et al. (2025). The asymmetric opening of HIV-1 Env by a potent CD4 mimetic enables anti-coreceptor binding site antibodies to mediate ADCC. Nat Commun 16, 10419. 10.1038/s41467-025-65866-x.

68. Scharf, L., Scheid, J.F., Lee, J.H., West, A.P., Jr., Chen, C., Gao, H., Gnanapragasam, P.N., Mares, R., Seaman, M.S., Ward, A.B., et al. (2014). Antibody 8ANC195 reveals a site of broad vulnerability on the HIV-1 envelope spike. Cell Rep 7, 785–795. 10.1016/j.celrep.2014.04.001.

69. Kong, R., Xu, K., Zhou, T., Acharya, P., Lemmin, T., Liu, K., Ozorowski, G., Soto, C., Taft, J.D., Bailer, R.T., et al. (2016). Fusion peptide of HIV-1 as a site of vulnerability to neutralizing antibody. Science 352, 828–833. 10.1126/science.aae0474.

70. Williams, L.D., Ofek, G., Schatzle, S., McDaniel, J.R., Lu, X., Nicely, N.I., Wu, L., Lougheed, C.S., Bradley, T., Louder, M.K., et al. (2017). Potent and broad HIV-neutralizing antibodies in memory B cells and plasma. Sci Immunol 2. 10.1126/sciimmunol.aal2200.

71. Krebs, S.J., Kwon, Y.D., Schramm, C.A., Law, W.H., Donofrio, G., Zhou, K.H., Gift, S., Dussupt, V., Georgiev, I.S., Schatzle, S., et al. (2019). Longitudinal Analysis Reveals Early Development of Three MPER-Directed Neutralizing Antibody Lineages from an HIV-1-Infected Individual. Immunity 50, 677–691 e613. 10.1016/j.immuni.2019.02.008.

72. Huang, J., Ofek, G., Laub, L., Louder, M.K., Doria-Rose, N.A., Longo, N.S., Imamichi, H., Bailer, R.T., Chakrabarti, B., Sharma, S.K., et al. (2012). Broad and potent neutralization of HIV-1 by a gp41-specific human antibody. Nature 491, 406–412. 10.1038/nature11544.

73. Croft, J.T., Do, H.N., Leaman, D.P., Lovendahl, K.N., Ralli-Jain, P., Chase, K.J., Chen, C., Prasad, V.M., Derdeyn, C.A., Zwick, M.B., et al. (2026). Structure of HIV-1 Env glycoprotein on virions reveals an alternative fusion subunit organization and native membrane coupling. bioRxiv. 10.64898/2026.01.09.698652.

74. Ruprecht, C.R., Krarup, A., Reynell, L., Mann, A.M., Brandenberg, O.F., Berlinger, L., Abela, I.A., Regoes, R.R., Gunthard, H.F., Rusert, P., and Trkola, A. (2011). MPER-specific antibodies induce gp120 shedding and irreversibly neutralize HIV-1. J Exp Med 208, 439–454. 10.1084/jem.20101907.

75. Foulkes, C., Friedrich, N., Ivan, B., Stiegeler, E., Magnus, C., Schmidt, D., Karakus, U., Weber, J., Gunthard, H.F., Pasin, C., et al. (2025). Assessing bnAb potency in the context of HIV-1 envelope conformational plasticity. PLoS Pathog 21, e1012825. 10.1371/journal.ppat.1012825.

76. Chatterjee, D., Niu, L., Medjahed, H., Ding, S., Benlarbi, M., Belanger, E., Prevost, J., Chen, H.C., Tolbert, W.D., Smith, A.B., 3rd, et al. (2025). A gp41 HR2 residue modulates the susceptibility of HIV-1 envelope glycoproteins to small molecule inhibitors targeting gp120. J Virol 99, e0226724. 10.1128/jvi.02267-24.

77. Bennett, A.L., and Henderson, R. (2021). HIV-1 Envelope Conformation, Allostery, and Dynamics. Viruses 13. 10.3390/v13050852.

78. Banach, B.B., Pletnev, S., Olia, A.S., Xu, K., Zhang, B., Rawi, R., Bylund, T., Doria-Rose, N.A., Nguyen, T.D., Fahad, A.S., et al. (2023). Antibody-directed evolution reveals a mechanism for enhanced neutralization at the HIV-1 fusion peptide site. Nat Commun 14, 7593. 10.1038/s41467-023-42098-5.

79. Kwon, Y.D., Georgiev, I.S., Ofek, G., Zhang, B., Asokan, M., Bailer, R.T., Bao, A., Caruso, W., Chen, X., Choe, M., et al. (2016). Optimization of the Solubility of HIV-1-Neutralizing Antibody 10E8 through Somatic Variation and Structure-Based Design. J Virol 90, 5899–5914. 10.1128/JVI.03246-15.

80. Parthasarathy, D., Pothula, K.R., Ratnapriya, S., Cervera Benet, H., Parsons, R., Huang, X., Sammour, S., Janowska, K., Harris, M., Sodroski, J., et al. (2024). Conformational flexibility of HIV-1 envelope glycoproteins modulates transmitted/founder sensitivity to broadly neutralizing antibodies. Nat Commun 15, 7334. 10.1038/s41467-024-51656-4.

81. Flemming, J., Wiesen, L., and Herschhorn, A. (2018). Conformation-Dependent Interactions Between HIV-1 Envelope Glycoproteins and Broadly Neutralizing Antibodies. AIDS Res Hum Retroviruses 34, 794–803. 10.1089/AID.2018.0102.

82. Poss, M., and Overbaugh, J. (1999). Variants from the diverse virus population identified at seroconversion of a clade A human immunodeficiency virus type 1-infected woman have distinct biological properties. J Virol 73, 5255–5264. 10.1128/JVI.73.7.5255-5264.1999.

83. Haddox, H.K., Dingens, A.S., Hilton, S.K., Overbaugh, J., and Bloom, J.D. (2018). Mapping mutational effects along the evolutionary landscape of HIV envelope. Elife 7. 10.7554/eLife.34420.

84. Kremer, J.R., Mastronarde, D.N., and McIntosh, J.R. (1996). Computer visualization of three-dimensional image data using IMOD. J Struct Biol 116, 71–76. 10.1006/jsbi.1996.0013.

85. Agulleiro, J.I., and Fernandez, J.J. (2015). Tomo3D 2.0--exploitation of advanced vector extensions (AVX) for 3D reconstruction. J Struct Biol 189, 147–152. 10.1016/j.jsb.2014.11.009.

