## Supplementary figures and images for "Distinct allosteric remodeling of HIV-1 Env dynamics on virions by gp41-directed antibodies reveals two modes of neutralization"

### Figure S1.tif

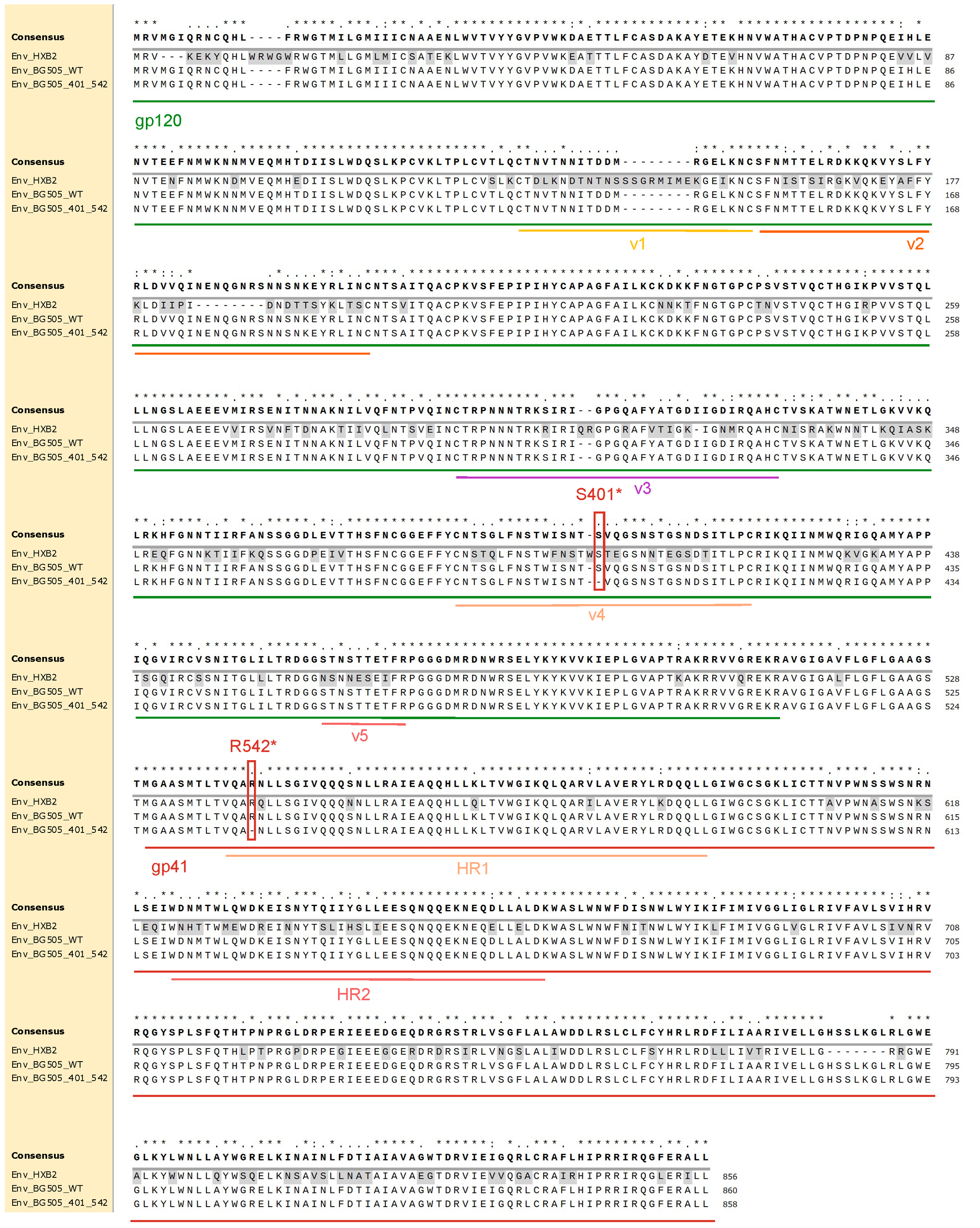

### Figure S2.tif

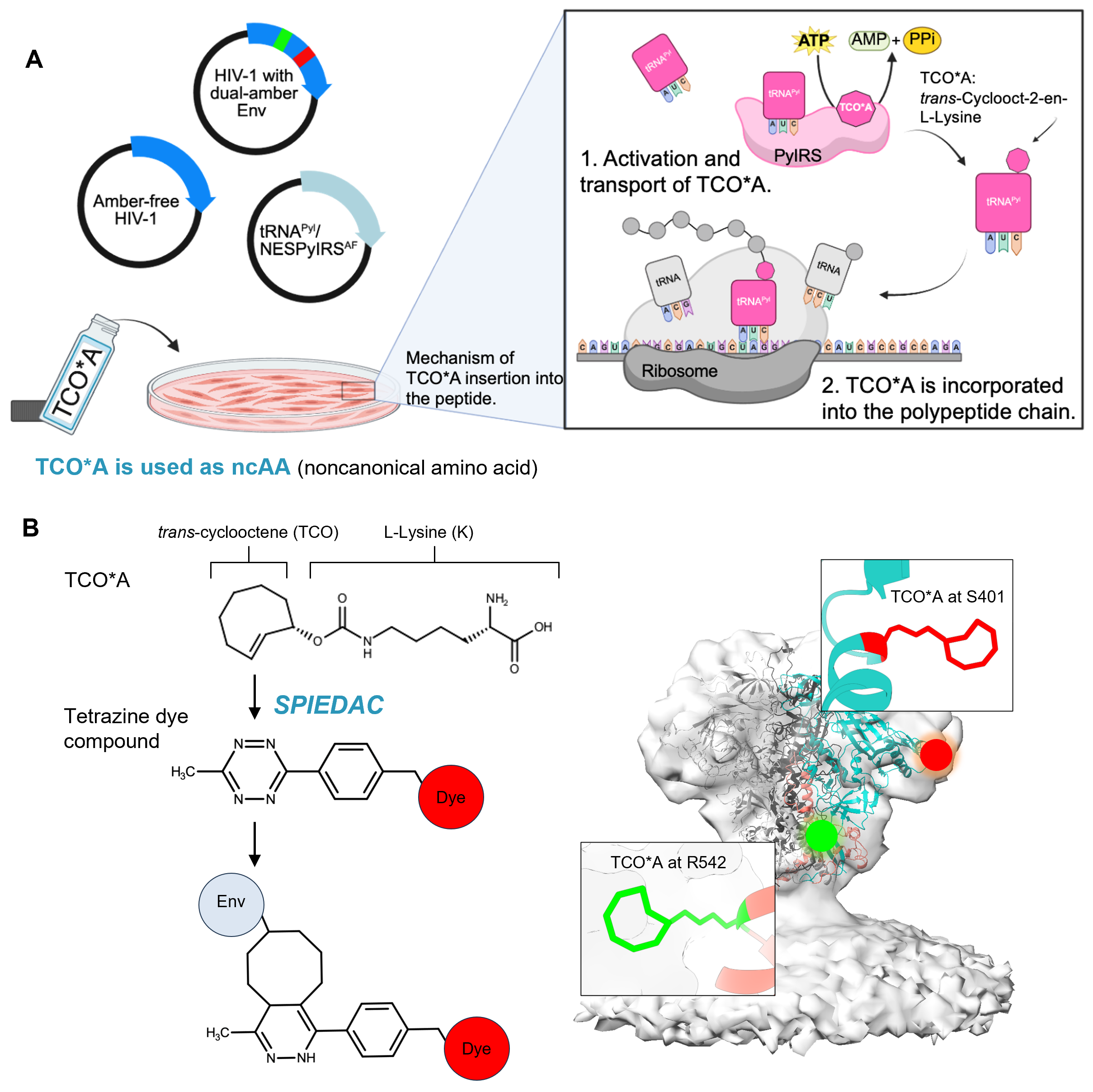

### Figure S3.tif

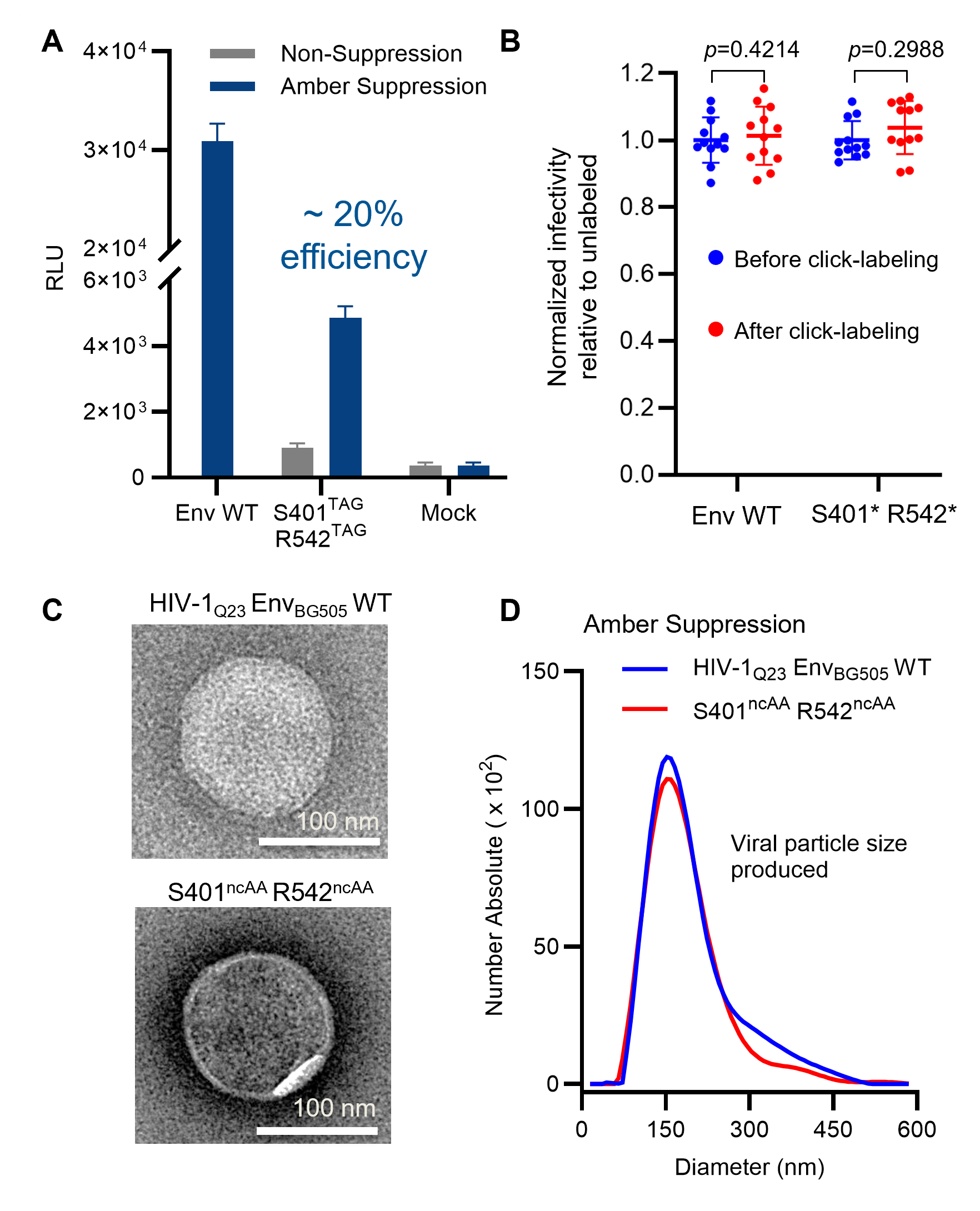

### Figure S4.tif

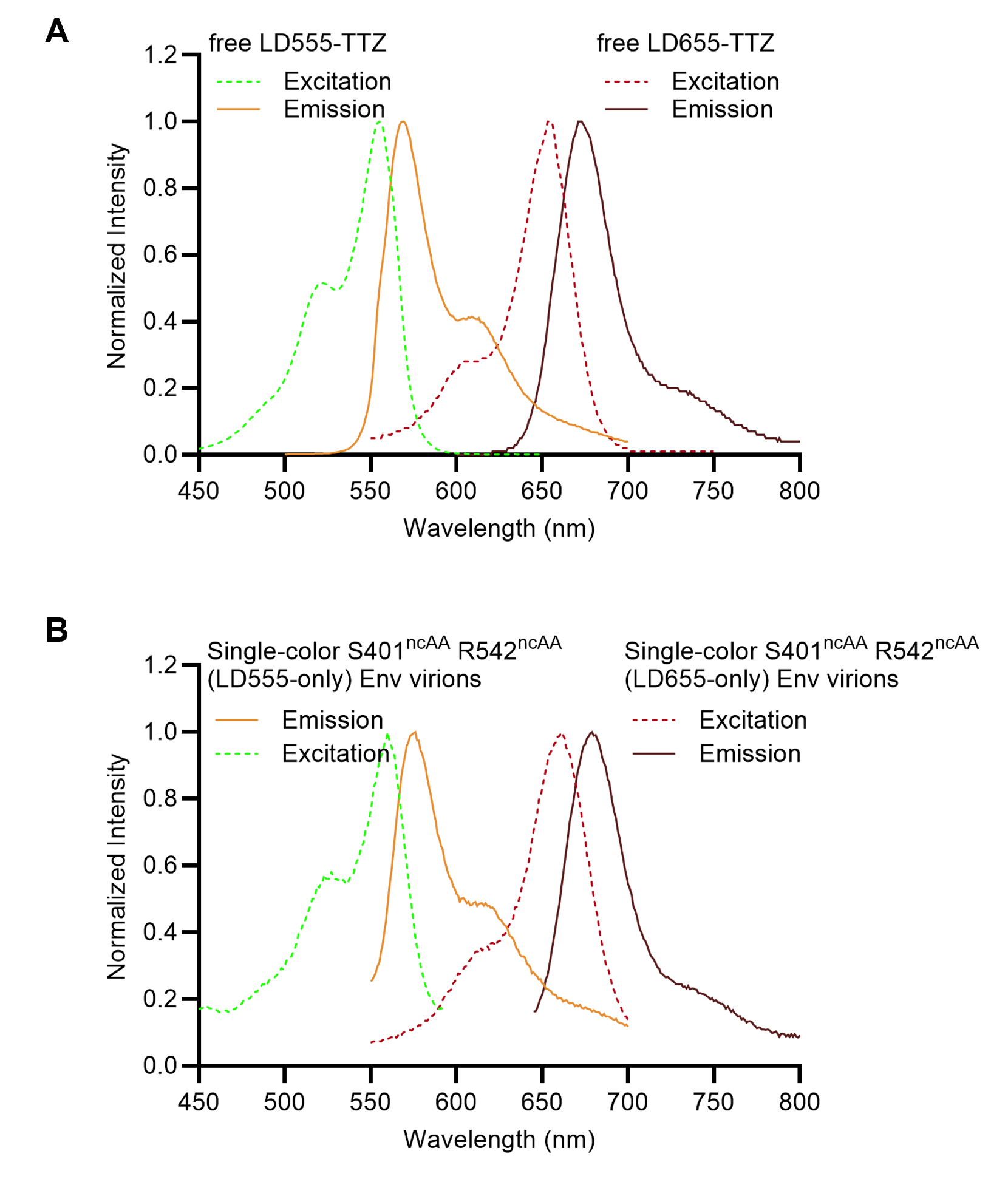

### Figure S5.tif

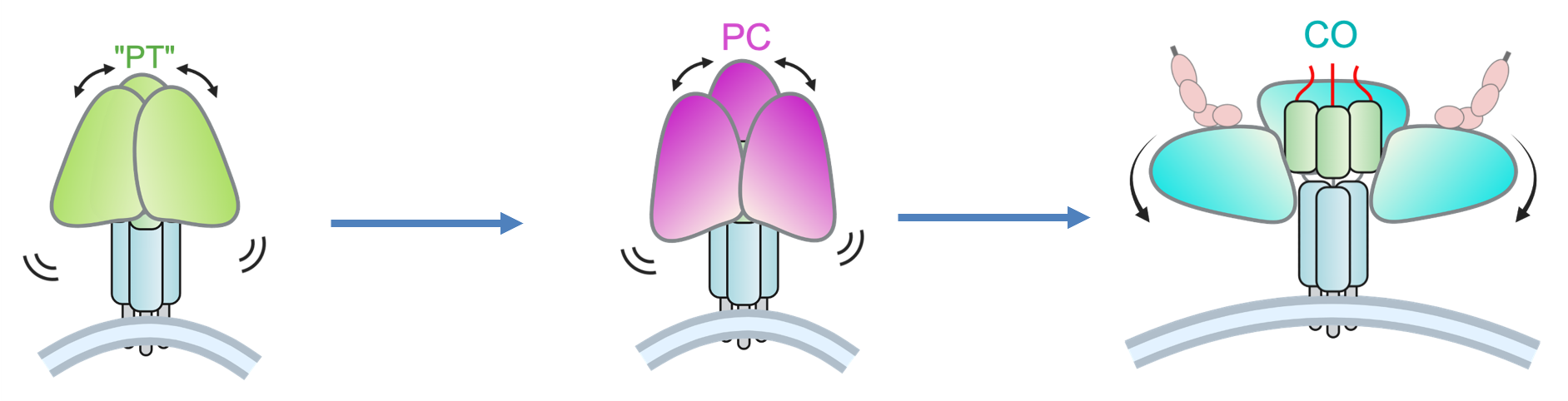

### Figure S6.tif

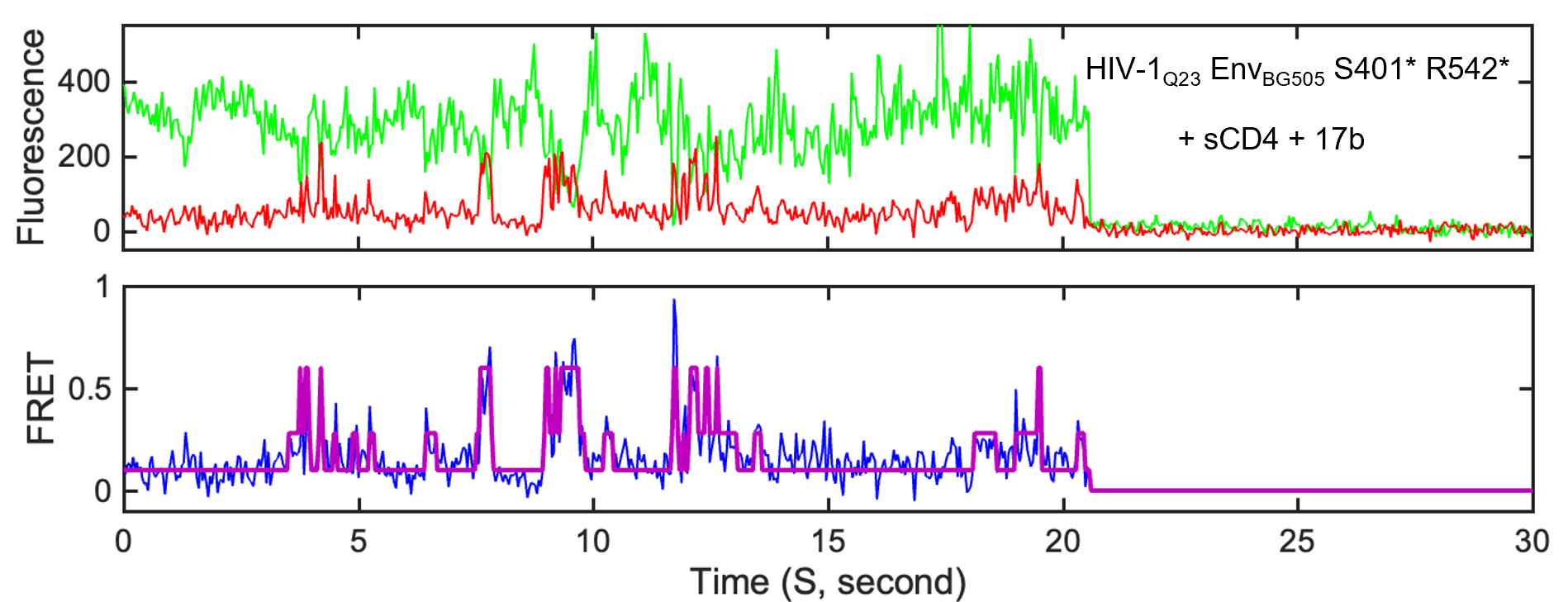

### Figure S7.tif

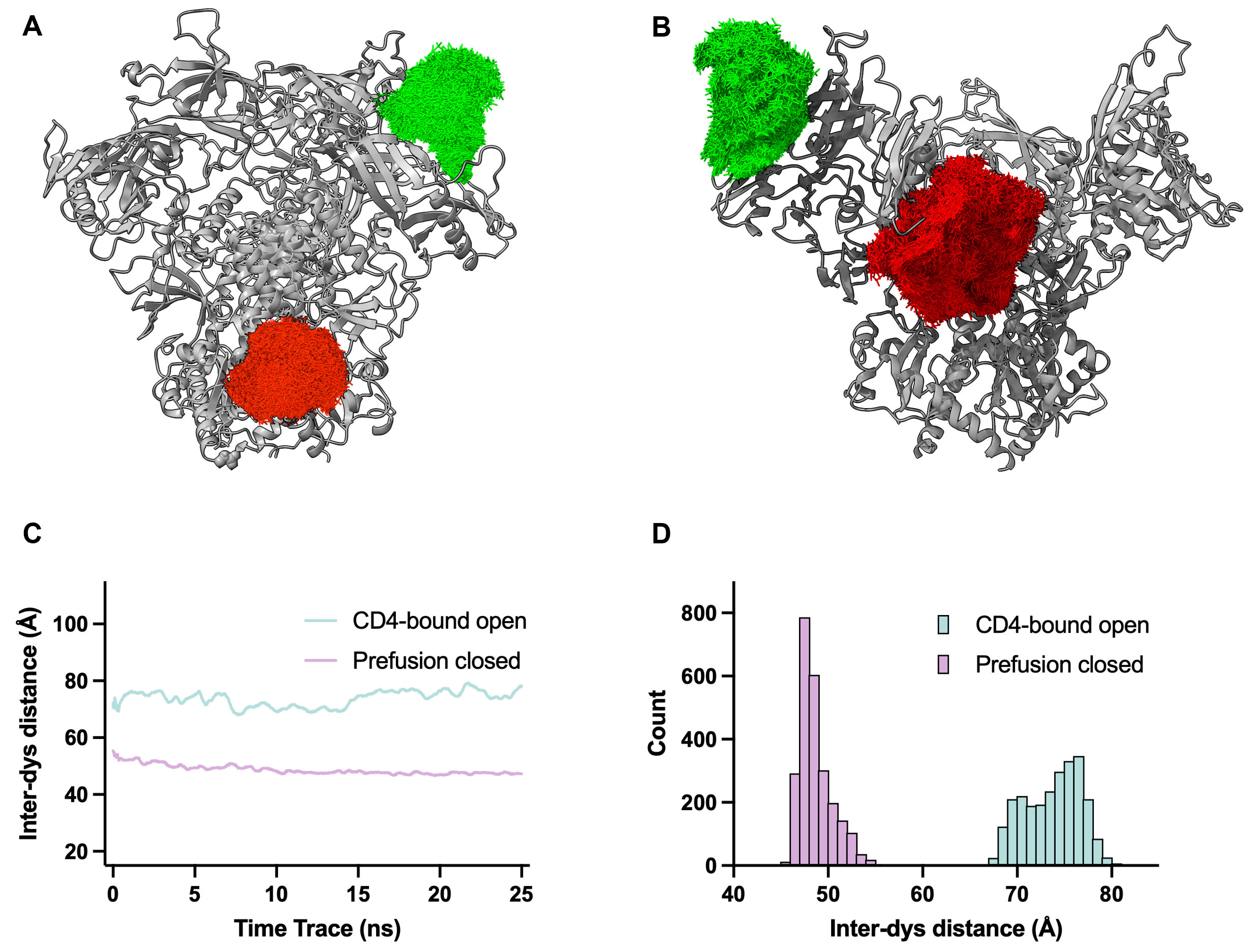

### Figure S8.tif

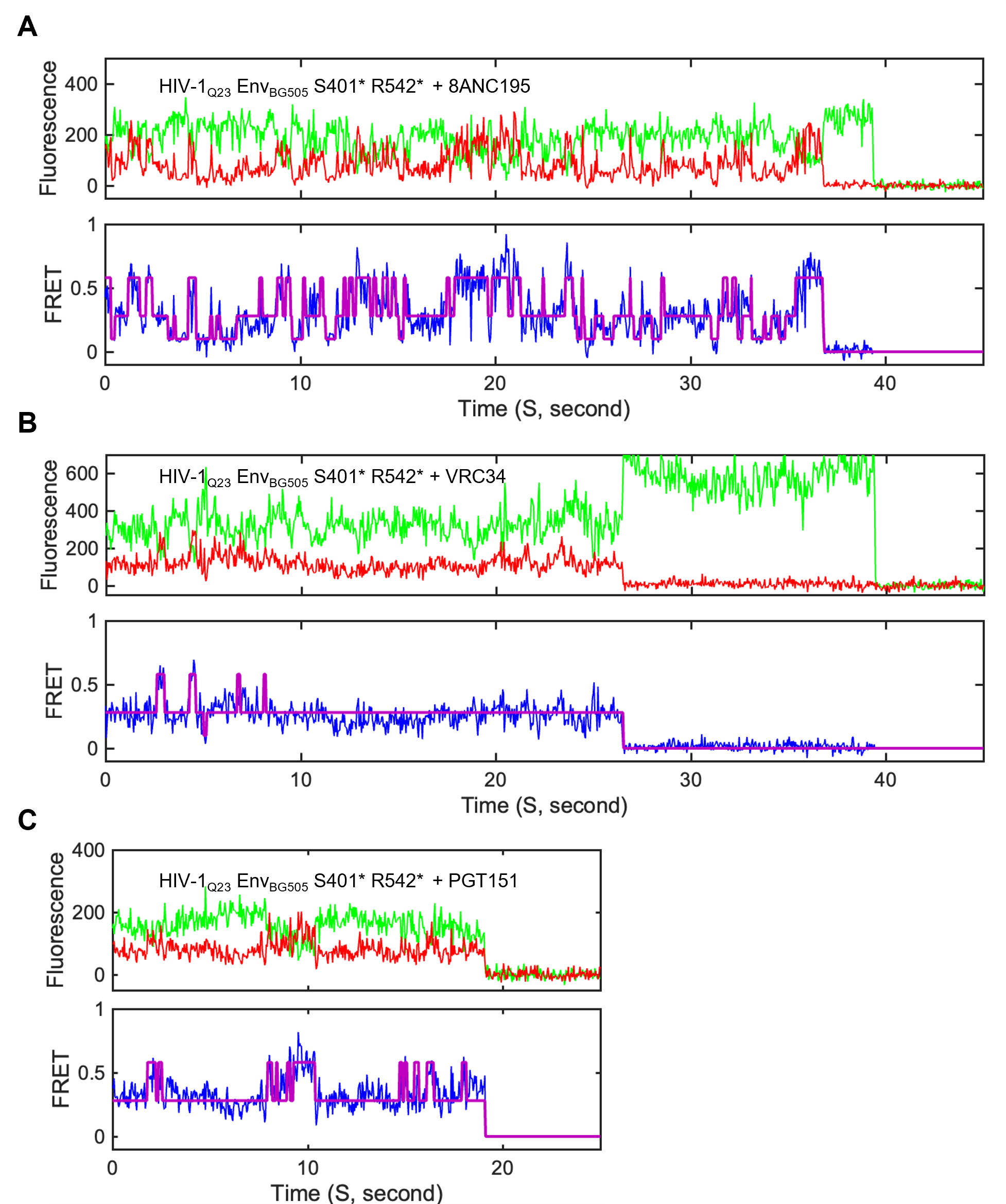

### Figure S9.tif

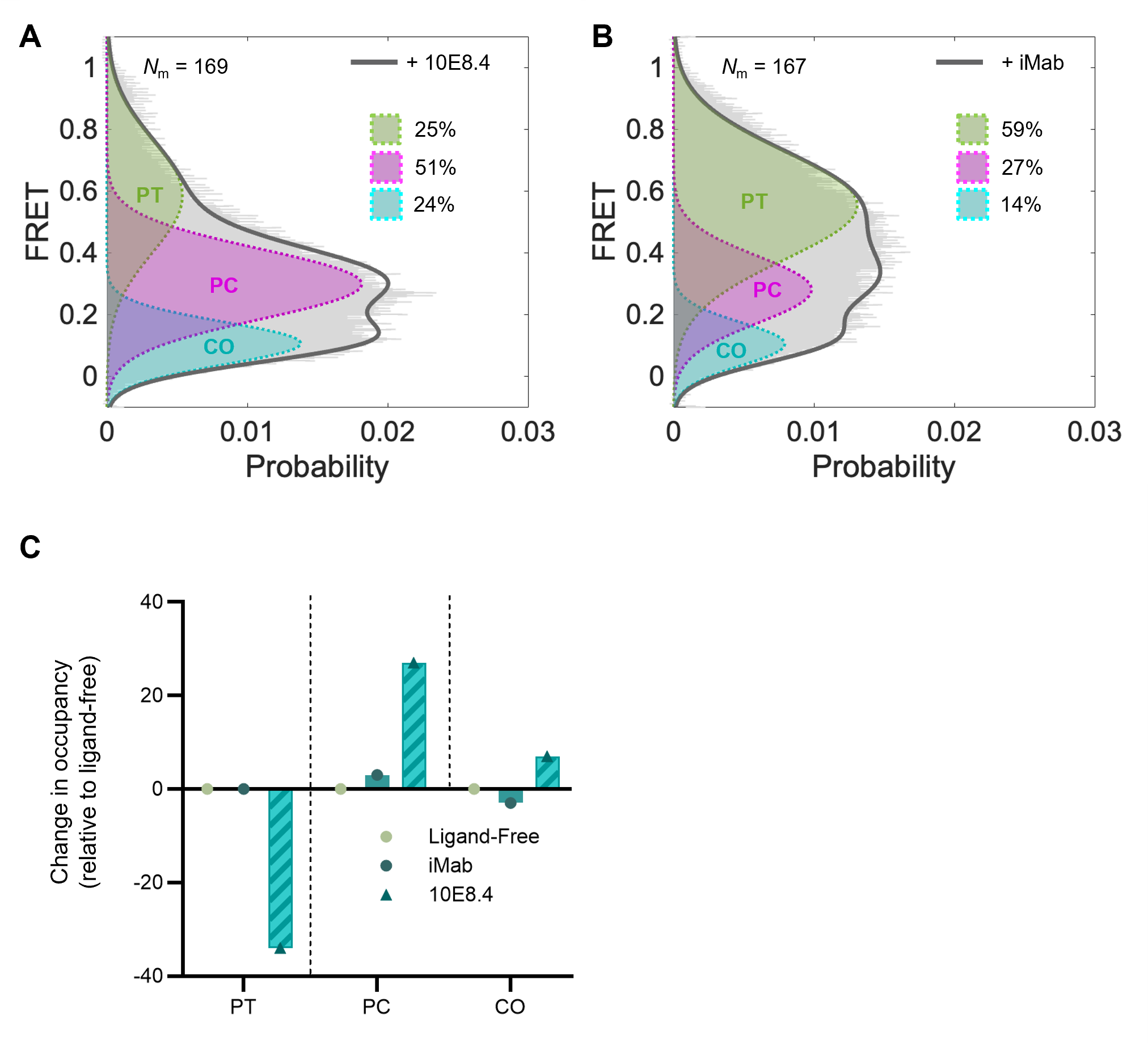

### Figure S10.tif

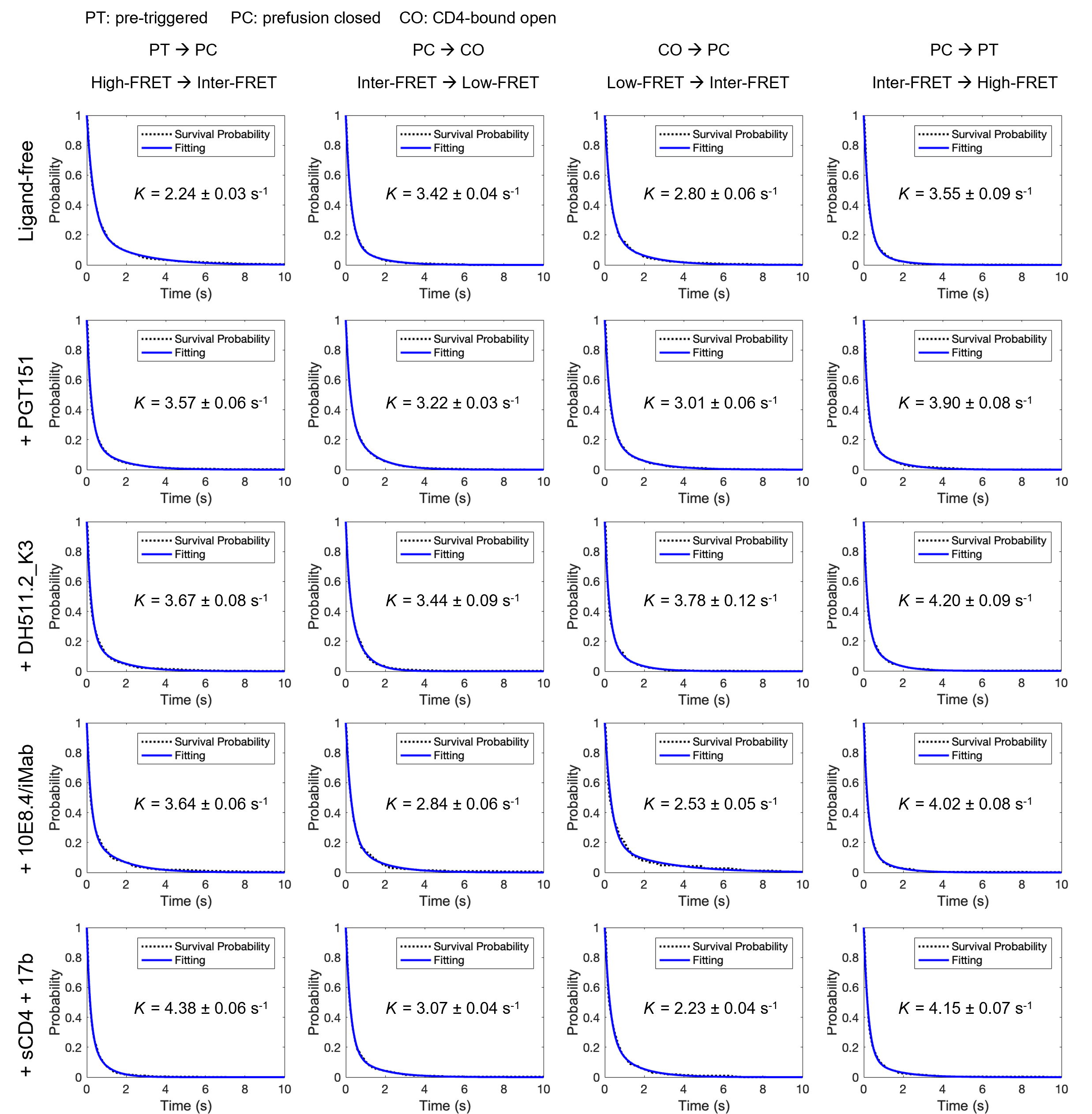
